# Complex nitrogen redox couplings control methane emissions from Arctic upland yedoma taliks

**DOI:** 10.1101/2025.02.09.637290

**Authors:** Oded Bergman, Katey Walter Anthony, E. Eliani-Russak, Orit Sivan

## Abstract

Yedoma-permafrost holds disproportionately large carbon and nitrogen pools, concentrated in icy, Pleistocene-aged silt deposits in the Arctic. Upon thaw, these undergo microbial mineralization, releasing greenhouse gases (GHGs) including carbon-dioxide (CO_2_), methane (CH_4_) and nitrous-oxide (N_2_O). Here we present combined geochemical data with microbial function and community dynamics from deep-talik soil boreholes in an unsaturated yedoma upland. Our results reveal significant in-situ spatio-temporal seasonal shifts in microbial functional, community composition and diversity within 7-m deep upland talik. In situ methanogenesis persisted in the soil talik throughout the year due to the permafrost thaw. In the winter methanotrophy was negligible within and above the methanogenic zone, leading to elevated CH_4_ emissions to the atmosphere. This is likely due to reduced microbial methanotrophic activity, associated with lower temperatures and nitrogen availability. During summer, at and above the anoxic methanogenic zone, nitrate/nitrite mediated anaerobic oxidation of methane (N-AOM) by ANME2d and the NC-10 phylum, together with aerobic methanotrophy near the soil surface, significantly attenuated CH_4_ emissions. Nitrous-oxide concentrations peaked at 10 cm (7.2 µM) and 105 cm (6.7 µM) and were associated with denitrification and N-AOM by *Methanoperedens* (ANME2d). In the summer only and within the top 1 m of soil, high expression of nitrogen related genes (narG, norB, amoA, Annamox, and Feammox) indicated active redox dynamics, potentially providing nitrogen species for N-AOM. The potential N_2_O emissions in summer may imply higher net GHGs emission from yedoma uplands as climate warming leads to longer summers and warmer soils in the future.

## Introduction

Permafrost soils cover about 24% of the northern hemisphere land surface. Approximately 25% of the permafrost soil carbon pool is concentrated in silty, ice-rich, Pleistocene-aged yedoma permafrost in Siberia and Alaska, which makes up 4%-7% of the permafrost region [1–3]. The carbon pool of terrestrial permafrost soils is estimated between 1330-1580 Pg [4], including 327–466 Pg in yedoma [2]. While less definitive, the nitrogen pool is estimated at 41.2 Pg at the upper 20 m of the yedoma domain [5].

The increase in air and ground temperatures leads to permafrost thaw and the formation of intra-permafrost zones of perennially thawed soil termed taliks [6–9]. Upon thaw, the immense carbon and nitrogen reservoirs may become available for microbial mineralization [10–12], and become a significant source of atmospheric CO_2_, CH_4_ and N_2_O emissions in yedoma landscapes as well as subarctic tundra, arctic peatlands, ponds, and lakes [5, 13–19]. methane and N_2_O are potent ozone-depleting greenhouse gases (GHGs) with a global warming potential roughly 300 and 33 times that of CO_2_ over a 100-year timescale [20].

Microbial communities differ significantly in composition and structure across various permafrost environments, presenting dominant microbial groups, such as *Proteobacteria* and *Actinobacteria* [21–26]. Concomitantly, community shifts have been related to permafrost thaw in-vivo and in incubation experiments [21, 27–31]. The widespread distribution of methanogenic and methanotrophic microbial communities in permafrost is also well documented, together with shifts in composition related to permafrost thaw and talik formation [21, 32–37]. Methanotrophy in permafrost sediments is performed by both anaerobic archaea (ANMEs) and aerobic bacteria (mainly Gammaproteobacteria) [38–42].

Anaerobic oxidation of CH_4_ (AOM) can be coupled to N-AOM. For example, AOM coupled to nitrate (NO_3_^-^) and nitrite (NO_2_^-^) reduction, has been shown by anaerobic bacteria including the NC10 phylum, in both freshwater and marine sediments [41, 43–47]. Under such settings, methanotrophy can co-occur alongside N_2_O production [48]. While N_2_O emissions were measured from thermokarst mounds in North-Siberia, a region of continuous-permafrost, where upland taliks have yet to develop [49], to our knowledge, the potential for N_2_O production in warmer regions, such as interior Alaska has never been explored. While N and C mobilization are interconnected through microbial and ecosystem processes, their dynamics are complex and require thorough investigation [31, 50].

This study focuses on the couplings between the CH_4_ and nitrogen cycles and the potential for N_2_O emissions in unsaturated yedoma upland taliks in interior Alaska. Our warmer study site, with advanced thaw, allows exploration of the potential carbon and nitrogen dynamics across the yedoma domain, as permafrost thaws and taliks develop in the future. Soil boreholes up to 7.1 m long were collected from the talik of an upland yedoma thermokarst field, informally named North Star Yedoma (NSY) (Fig. 1). Geophysical surveys indicate a 5 to 9 m thick talik across NSY; this thaw, which led to the still-ongoing melting of ground ice wedges, resulted in regularly spaced thermokarst mounds that characterize polygonal ground in yedoma regions [51, 52]. We structured our analysis around three main axes of comparison to elucidate key patterns and processes. In Axis 1 we explore seasonality differences between summer and winter soils in the surface 3 m. In Axis 2 we assessed shallow vs. deep soil layers within a single winter 7-m deep borehole, analyzing the full talik profile down to the top of permafrost. Finally, in Axis 3 we examined field-scale differences in elevation corresponding to shallower (mid-elevation site) vs. deeper (high-elevation site) surface aerobic zones.

To achieve our goals, we utilized a combination of soil geochemical methods, together with advanced molecular techniques.

**Figure 1.**
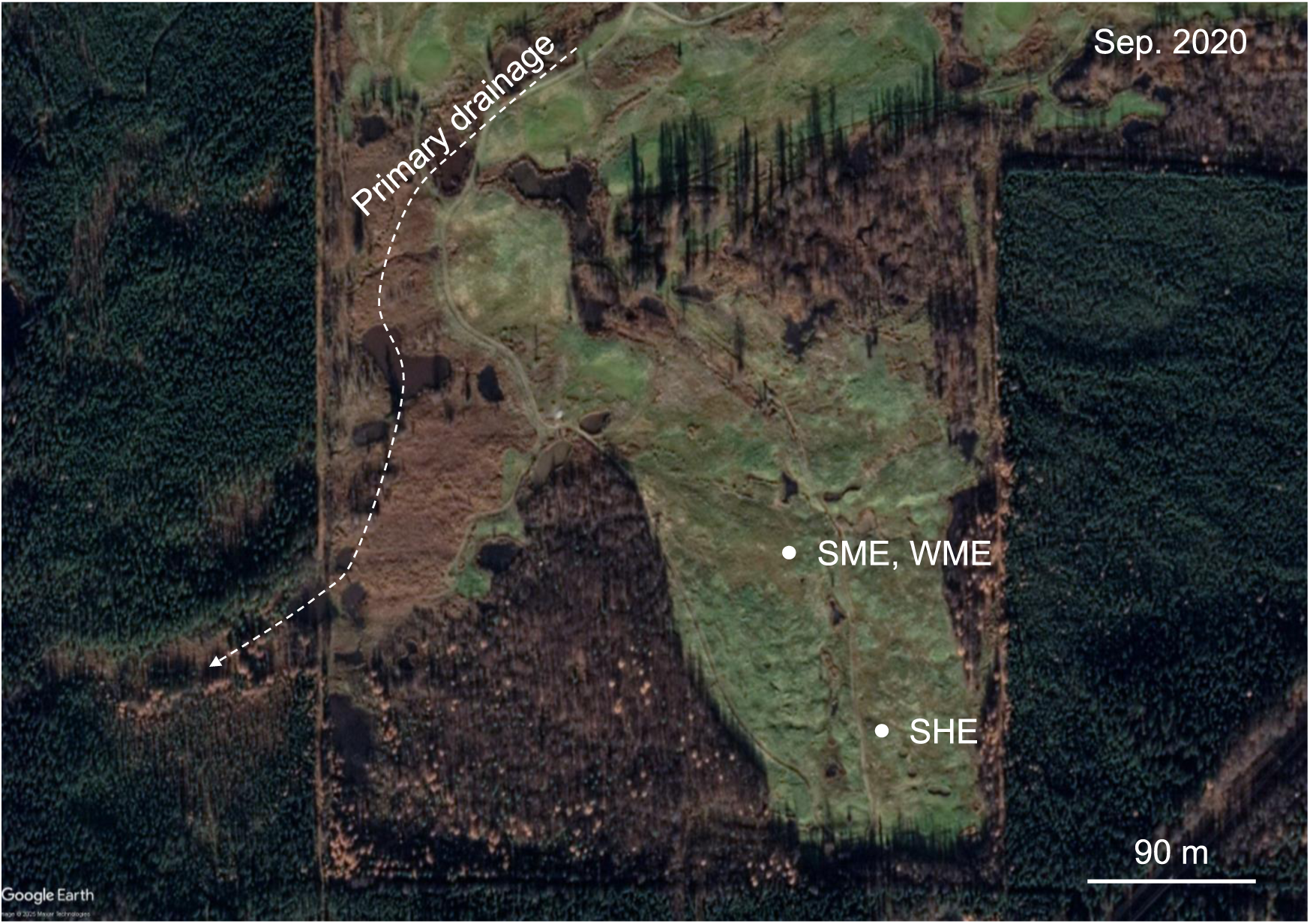
Map of study site. The study site was located at North Star Yedoma (NSY) (Latitude: 64.8939, Longitude: −147.6373), located 7 km north west of Fairbanks, Alaska. Three boreholes were drilled and sampled. During summer (September 15, 2021) two shallow boreholes were excavated: SME = Summer Mid Elevation (3.2 m, borehole ID: BH1), SHE = Summer High Elevation (2.3 m, BH2). One deep borehole was drilled during winter (March 18, 2023): WME = Winter Mid Elevation (7.25 m, BH6). The upland yedoma field is characterized by thermokarst mounds and a 5 to 9 m deep talik. SHE is at about 623 feet elevation, SME is about 609 feet elevation, and the drainage path at the valley bottom is 585 feet. BH1 and BH6 were excavated 60 cm apart.

## Materials and methods

### Study site

The NSY study site is located in interior Alaska, seven kilometers north-west of Fairbanks. Originally a mature black spruce forest, this large open field contains thermokarst features, formed following an anthropogenic disturbance. Subsequently, intense thermokarst-mound were developed in the eastern half of the field, which was latter seeded with turf grasses for establishment of a rugged golf course. The western side was undisturbed, allowing mound development and natural forest succession. Over the last 20 years, ground-ice melt and thermokarst subsidence continued across the whole field resulting in the mound-ridden surface at NSY today. Borehole data of NSY surrounding area indicate organic-rich silts extending over 40 m below ground. The study site is extensively described in Walter Anthony et al., 2024 [37], including vegetation, soil description, disturbance history, exc.

### Sample collection, preparation and physicochemical analysis

Soil samples were collected during summer by drilling two boreholes near the thermokarst mounds at two locations, on September 15, 2021 (Fig. 1). The first was located at mid elevation (BH1, 3.2 m depth) and the second near the highest elevation in the field (BH2, 2.3 m). Boreholes were drilled using a gas-powered AMS frozen soil auger with 5 cm of internal (core) diameter. On March 18, 2023, a winter core (BH6, 7.25 m) was drilled 60 cm away from BH1, using a Talon Drill system. Subsamples from each core, not volumetrically controlled, were collected at 10 to 15-cm intervals, placed in ziploc bags, and transferred to the freezer. Additional description can be found at Walter Anthony et al., 2024 [37].

### Environmental parameters

The following physicochemical parameters were measured: Volumetric Water Content (VWC) Gravimetric Water Content (GWC), CH_4_ and CO_2_ concentrations, organic Carbon (C_org_, %), organic Nitrogen (N_org_, %), δ^15^N (‰), δ^13^C (‰). The full, detailed description of the methods used is presented in Walter Anthony 2024. Briefly, GWC was determined as the weight loss after drying at 105 °C, expressed as a percentage of wet weight. VWC was calculated as the product of gravimetric moisture content and dry density. Where dry density was not measured, site-specific or depth-dependent relationships were used. Dissolved CH_4_ and CO_2_ concentrations were determined using 3 ml soil plug samples placed in 20 mL vials containing 5M NaCl, sealed with butyl rubber stoppers, and stored upside down to prevent gas leakage. Gas concentrations (CO_2_, CH_4_ and N_2_O) were analyzed using a Focus Gas Chromatograph (GC) system (Thermoscientific, Germany) equipped with a flame ionization detector and shinCarbon ST packed column (Restek, USA). C_org_ and N_org_ were measured using an elemental analyzer (Costech ECS4010) coupled to a Finnigan DeltaPlus XP Isotope Ratio Mass Spectrometer (Thermo Scientific). Samples were acidified with 31.45% HCl to remove inorganic carbon, rinsed, and dried prior to analysis. C_org_ and N_org_ concentrations were reported in weight percentage (wt%). δ^13^C and δ^15^N isotopic compositions were measured simultaneously during the analysis and are expressed in per mil (‰) relative to V-PDB and ambient air, respectively. For the full methodological details, see Walter Anthony (2024) [37].

### PCA analysis

We sought to characterize the differences in various environmental conditions related to both sampling layer and sampling season (mixis vs. stratified periods). We performed an initial PCR analysis (n=27) on all samples from BH1 (n=10), BH2 (n=6) and BH6 (n=11) (Figs. S1A-S1B and Tables S1). We initially included 9 environmental parameters: VWC, GWC, CH_4_ and CO_2_ concentrations, C_org_, N_org_, C_org_/N_org_ ration, δ^15^N and δ^13^C. After omitting the variables showing strong cross-correlation, or low loading scores (Tables S2-S3), a final PCA was performed with 4 environmental parameters: GWC, CH_4_ and CO_2_ concentrations, and C_org_. Prior to PCA analysis all samples were centered and scaled (log+1 values). To maintain the three main axes of comparison, from hereafter we have denoted the sampled cores by season and elevation, as follows: BH1 = SME: Summer Mid Elevation (n=10), BH2 = SHE: Summer High Elevation (n=6), BH6 = WME: Winter Mid Elevation (n=11). The WME samples were comprised of shallow (<3 m, n=5) and deep (<3 m, n=6) samples. Summer sampling refers to cores taken on the September 15, 2021 and winter on March 18, 2023.

### DNA extraction

Soil samples from BH1 (23, 56, 68, 86, 106, 166, 186, 259, 295 and 305 cm), BH2(8, 38, 84, 146, 195 and 225 cm) and BH6 (50, 100, 162, 198, 285, 345, 535, 564, 655, 695 and 711 cm) were taken for molecular microbial analysis. DNA was extracted from approximately 0.25 gr soil, using the PowerSoil™ DNA Isolation Kit (QIAGEN, Hilden, Germany; formerly MoBio, CA, USA), according to the manufacturer’s instructions, with the following modifications: After lysis buffer addition, samples were incubated at 65 °C for 10 minutes to enhance lysis. The elution buffer was pre-warmed at 70 °C for 5 minutes to improve DNA recovery. DNA extracts were subsequently stored at −80 °C until use.

### Real time PCR (qPCR)

qPCR was performed using a CFX Duet qPCR instrument (Bio-Rad, USA) and analyzed with the CFX Maestro software. Reactions targeted genes associated with the following processes: methanogenesis (mcrA), methane oxidation (pmoA), anammox (hzsB), Feammox (16S gene for *Acidimicrobiaceae* bacterium A6), aerobic ammonium oxidation (amoA, bacterial and archaeal), NC10 phylum (16S gene), and denitrification (narG, nirK, and norB). Target genes primer names and concentrations (mM), and accession numbers for gBlocks (Integrated DNA Technology, UK) used to construct the standard curves are provided in Table S11. gBlocks were re-suspended in Tris-EDTA (10 mM Tris-HCl, 1 mM disodium EDTA, pH 8), following the manufacturer’s instructions, adjusted to a final concentration of 10 ng/µL, and stored at −20 °C until use. Linearized plasmids were used for the narG and norB genes as standards. PCR products of the correct size were gel-extracted using the Nucleospin PCR Clean-up Kit (Macherey-Nagel, Germany) and cloned into E. coli JM109 cells using the pGEM-T Easy vector system (Promega, USA). Clones were grown in TYP broth and plasmids were extracted using the Nucleospin Plasmid EasyPure kit (Macherey-Nagel, Germany). Sequencing of plasmids was conducted at HyLabs (Israel) using T7-SP6 primers, and product identities were confirmed through BLAST searches. Plasmids containing target products were linearized with the SacI restriction enzyme, purified, and used for standard curve generation through serial dilutions to determine gene concentrations. The qPCR reactions were performed using the Fast SYBR Green Master Mix (Applied Biosystems, USA). The PCR reaction were carried out in a volume of 20 μL, containing 12.5 μL master Mix, primer concentration (between 0.15-0.5 mM) and 2 μL template DNA. PCR conditions included an activation step at 95 °C for 20 seconds, followed by 40 cycles of denaturation at 95 °C for 3 seconds and annealing/extension at 60 °C for 30 seconds. During initial calibration, reactions were visualized on 1.5% agarose gels to confirm specificity and amplification of the correct band size. Melt curves were generated for all the reactions, to assess specificity, confirming single amplicon by detecting a single distinct melting temperature peak.

### 16S rRNA gene V4 amplicon-sequencing

We performed 16S gene amplicon-based sequencing, using the modified primer pair with consensus sequences CS1_515F (ACACTGACGACATGGTTCTACAGTGCCAGCMGCCGCGGTAA) and CS2_806R (TACGGTAGCAGAGACTTGGTCTGGACTACHVGGGTWTCTAAT) (Sigma-Aldridge, Israel) [53]. The initial PCR was carried out in 25 μL reactions containing 12.5 μL of KAPA HiFi HotStart ReadyMix (KAPA Biosystems, Wilmington, WA, USA) and 0.75 μL of forward and reverse primers at a final concentration of 300 nM each. PCR conditions included an initial denaturation at 95 °C for 3 minutes, followed by 30 cycles of 98 °C for 20 seconds, 60 °C for 15 seconds, and 72 °C for 30 seconds. PCR products were visualized on a 2% agarose gel to assess band intensity. Samples were pooled and purified using calibrated Ampure XP beads before being used for library preparation. PCR visualization, purification, library preparation, and sequencing (2 × 250 bp paired-end reads) were conducted on an Illumina MiSeq (at HyLabs, Israel). Demultiplexing of paired-end reads and subsequent analyses were performed using QIIME2 (v2020.11) [Bolyen2019]. Sequencing quality was assessed using the q2-demux plugin, followed by chimera detection and merging of reads into Amplicon Sequence Variants (ASVs) using the q2-dada2 plugin[54]. ASVs were defined by clustering at 100% similarity [55] to account for length variations. Taxonomy was assigned using the SILVA 138 QIIME release database clustered at 99% similarity [56]. The classifier was trained with the extract-reads and fit-classifier-naive-bayes methods via the q2-feature-classifier plugin [57], and ASV classification was performed using the classify-sklearn method (ver. 0.23.1) [58]. The q2-diversity plugin was used to generate rarefaction curves at varying depths, to confirm ASVs reached a plateau and to justify the chosen rarifying depth (Fig. S2). Diversity analysis was also performed using the q2-diversity plugin, at a rarifying depth of 16,159 ASVs. Alpha (Shannon’s entropy, Pielou’s evenness index and Faith’s PD index) and beta (jaccard and bray-curtis) metrics were constructed and presented in Supplementary Results section 1.2. Venn diagrams were constructed based on the rarefied ASVs tables, obtained from the q2-diversity plugin. Downstream analyses were conducted in R using the phyloseq [59] and ggplot2 [60] packages. Based on the 16S rRNA data, we performed a pathway prediction analysis using the PICRUSt2’s (Phylogenetic Investigation of Communities by Reconstruction of Unobserved States) package [61]. The Kyoto Encyclopedia of Genes and Genomes (KEGG) database [62] was used to identify Orthology (KO) groups and modules (MO), related to the CH_4_ and nitrogen cycles (Supplementary data 6-7). Since 16S rRNA gene copy number variation introduces bias in relative abundance estimates, this bias propagates into functional predictions generated by PICRUSt2 [63]. To correct for this, we adjusted PICRUSt2 predictions using qPCR-derived gene abundances, providing a more accurate representation of functional potential in the microbial community.

Generated raw sequences reads were deposited in to the European Nucleotide Archive (ENA), at the EMBL European Bioinformatics Institute (EMBL-EBI) Database (https://www.ebi.ac.uk/ena/browser/home). BioProject accession number PRJEB59938. Additional data are available under the Supplementary Information and Supplementary Data sections.

### Statistical analysis

Statistical analyses were performed using the R (v. 4.0.3) rstatix package and the QIIME2 (v. 2020.11) software. For the PCA analysis, variables were log transformed, scaled and centered to account for variations in magnitude and units. Statistical analysis for all microbiome diversity analysis was done using the alpha-group-significance (Kruskal– Wallis tests) and beta-group-significance (PERMANOVA, PERMDISP, and ADONIS with 999 permutations). p-values were adjusted according to the Benjamini–Hochberg FDR correction. All statistical tests were two-sided. p-value was considered significant if < 0.05.

## Results and Discussion

### Axis 1 - Seasonal shifts between summer and winter

#### Unsaturated-upland yedoma soils represent a warmer more advanced state of talik formation

During summer, oxic conditions persisted down to 50 cm depth and temperatures gradually declined, from ∼9 °C (at 10 cm depth) to ∼5.5 °C (at 250 cm).. In winter, soil-surface (10 cm) temperature neared the freezing point (−0.42 °C), gradually elevating to 1.09 °C (250 cm). With soil-surface freezing, anaerobic conditions at the top 15 cm (−1.22%), turned to semi-aerated at 50 cm. Year-round anaerobic conditions were measured at 100 cm and 190 cm (∼-1.3%) (data taken from Walter_Anthony 2024 [37]. Our previous study, which included additional data, including modeling, remote sensing, geophysics, and field observations, showed that NSY represents a more advanced state of permafrost thaw and talik formation among upland yedoma landscapes, providing an opportunity to study these projected global warming effects [37]. Other studies on upland yedoma permafrost thaw and GHG generation focused mainly on the coldest permafrost regions of North Siberia (continuous permafrost), targeting near surface exposures and the active layer [11, 49, 64–67]. To the best of our knowledge, this study is the first to investigate the microbial community dynamics and function related to the CH_4_ and nitrogen cycles, in well-developed upland yedoma taliks.

### Seasonal changes in physico-chemical parameters

We characterized all sampled environments based on several physico-chemical parameters and performed a PCA analysis (PC1-PC2: 47.6-31.1%, n=27, Fig. 2A-B and Table S1). Up to ∼3 m depth, summer (SME) and winter (WME) samples grouped separately, with substantial variability withing each group. Summer samples, portrayed higher GWC, C_org_ and CO_2_ concentrations (Fig. 2B and Table S1 and S4).

**Figure 2.**
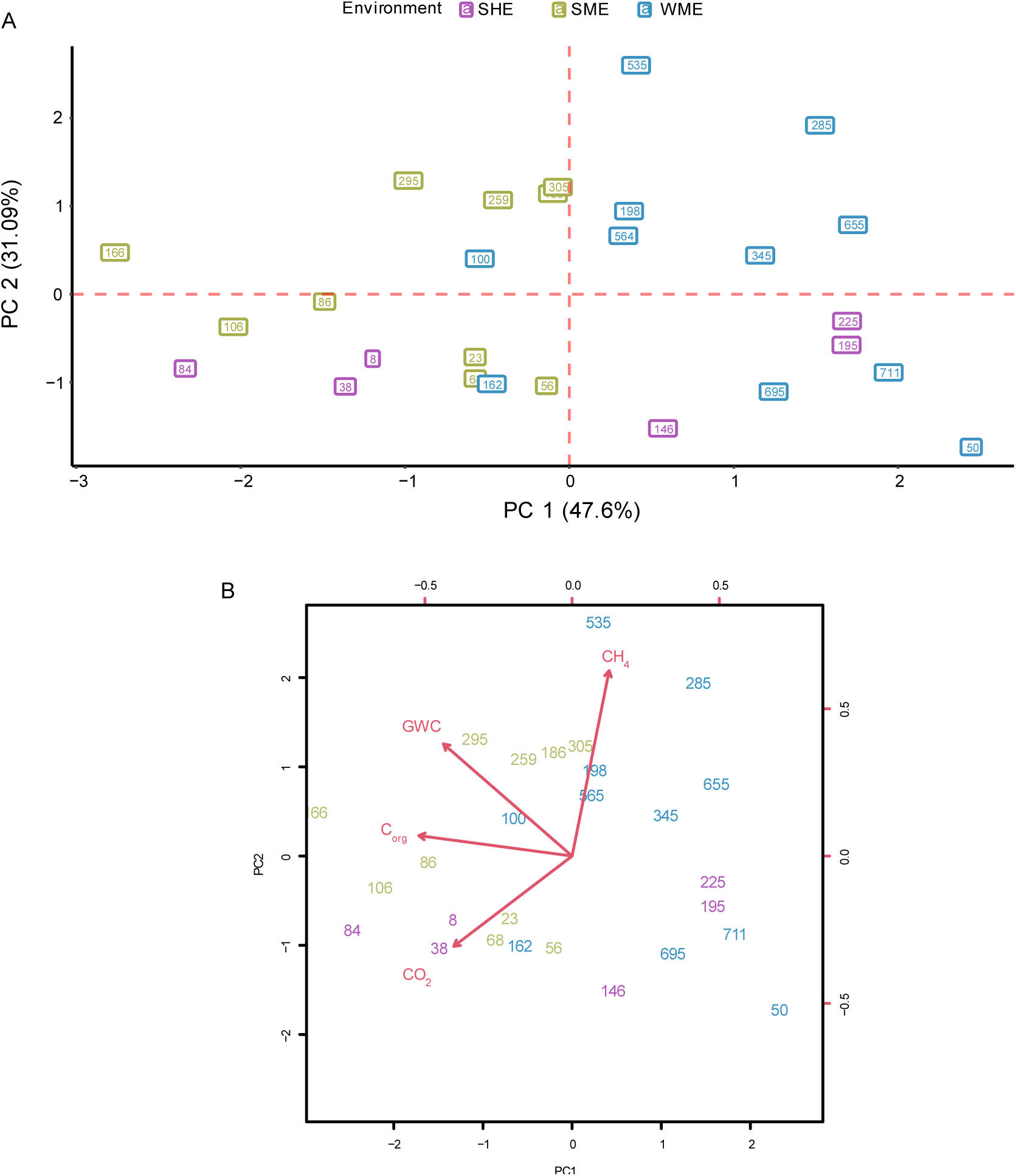
PCA analysis of selected environmental parameters at the NSY study site. (A) PCA analysis (n=27) of samples collected from BH1 (n=10), BH2 (n=6) and BH6 (n=11) was performed on samples collected during summer (BH1 and BH2, September 15, 2021) and winter (BH6, March 18, 2023). The final PCA included four environmental parameters (Table S1): Gravimetric Water Content (GWC), CH_4_ and CO_2_ concentrations, and organic Carbon (C_org_, %). To maintain the three main axes of comparison, we have denoted the sampled cores by season and elevation, as follows: SME = Summer Mid Elevation (borehole ID: BH1, n=10, olive-green), SHE = Summer High Elevation (BH2, n=6, purple), WME = Winter Mid Elevation (BH6, n=11, blue). Summer sampling refers to cores taken on the September 15, 2021 and winter on March 18, 2023. This final PCA was based on an initial PCA analysis (Fig. S1) that included nine environmental parameters. The variables with high collinearity or low loading scores were excluded from the final analysis. For additional details see Materials and Methos. **(B)** Loading scores, indicating the importance of tested environmental variables related to PC 1 and PC 2.

Methane concentrations were depleted close to the surface, increasing with depth and peaking between 185 to 295 cm (Fig. 3 and Table S1). During winter CH_4_ was present up to the soil-surface, peaking at greater depths (285-535 cm, elaborated in Axis 2 comparison below). Two peaks in N_2_O concentrations were found only in the summer profile (10 cm (7.2 µM) and 105 (6.7 µmole/L), Fig. 3); nitrous oxide was below detection levels in the winter core. Similar to previous reports, C_org_/N_org_ ratios were low in both seasons (SME: 13±0.45, WME: 11.42±0.46; Table S1), supporting the existence of a high N stock [49, 68] and potential increases in N_2_O production [11]. The counter behavior of CH_4_ and N_2_O raises significant concerns, as N_2_O is about nine times more efficient in trapping heat than CH_4_ [69]. During AOM by *Methylomirabilis Oxifera*, nitrite is reduced and used as an electron acceptor. By NO_2_^-^ labeling experiment Ettwig et al., (2010) [43] found that while most of the culture reduced NO_2_^-^ to N2, 7% was reduced to N_2_O by other denitrifiers. A similar ratio during denitrification in favor of N_2_O was also observed in thawed loamy soils [70]. From mass balance calculation, for 30 CH_4_ molecules that are anaerobically oxidized, one molecule of N_2_O is released. This will reduce CH_4_ sink of AOM by as much as 30% (as N_2_O is nine times more potent then CH_4_ as a GHG).

**Figure 3.**
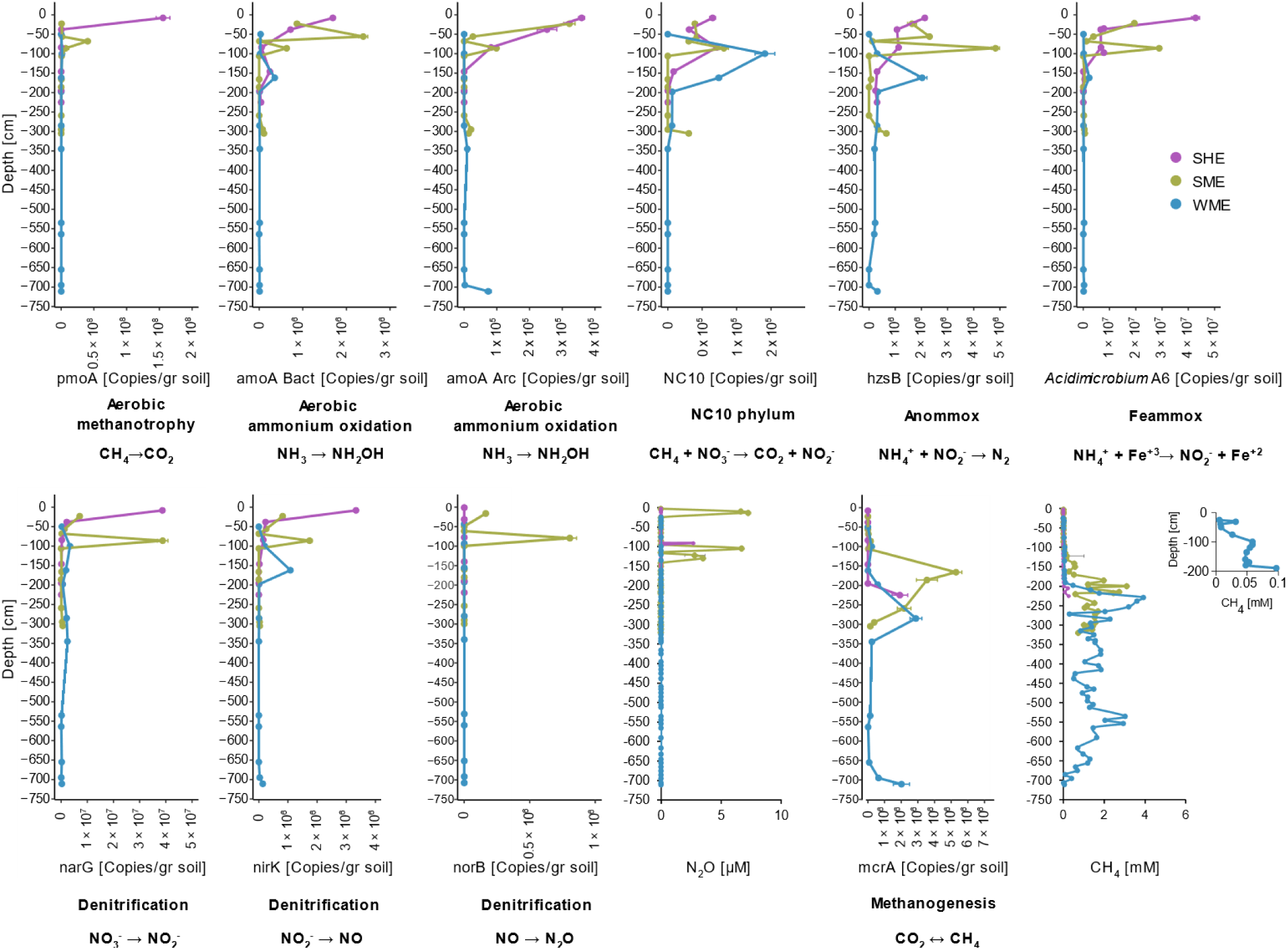
Absolute abundance depth profiles of genes related to the nitrogen and CH_4_ cycles. Quantification of absolute abundances (n=27 samples) was performed using qPCR with primers targeting functional genes related to the nitrogen and CH_4_ cycles (see Table S11 for a detailed list of genes and primers). For the Feammox reaction and NC10 phylum, functional genes are not available in the literature, and primers targeting regions of the 16S rRNA gene were used. For all sampled depths in the three cores, the absolute abundance of each gene is presented (copies/gr soil, mean±SE). For each gene, the related function and main reaction (main products, unbalanced) are indicated. For the mcrA gene, the double arrow symbol (↔) represents potential forward and reverse methanogenesis. nitrous oxide and CH_4_ concentrations are also presented. SME = Summer Mid Elevation (borehole ID: BH1, n=10, olive-green), SHE = Summer High Elevation (BH2, n=6, purple), WME = Winter Mid Elevation (BH6, n=11, blue). Summer sampling refers to cores taken on September 15, 2021 and winter on March 18, 2023.

### Seasonal shifts in methanogenesis and methanotrophy

Although both environments acted as a net CH_4_ source, highest emissions were measured during winter [37]. Winter CH4 emissions were 3 times higher than summer (165.8 ± 4.0 vs 55.7 ± 2.3 mg CH_4_ m^−2^ d^−1^, mean ± SEM), with the difference between the two attributed to methanotrophy. Winter CH_4_ peaks were observed at deeper depths, with elevated concentrations measured closer to the soil surface. Corresponding patterns were noted in mcrA (methanogenesis) and pmoA (methanotrophy) absolute gene expression. While higher mcrA levels were measured in summer across most anaerobic depths, peaking at 166 cm (Fig. 3 and Table S5), winter expression shifted to deeper depths.

The better understand these differences in gene expression, we assessed the microbial community composition. The microbial community in all tested environments was diverse, with most genera present at low relative abundance (<1%, Supplementary Data 1 and 2). The prevalent phyla were consistent with previous permafrost studies [21–26]. Microbial diversity was higher in summer, with about five times more unique archaeal and bacterial Amplicon Sequence Variants (ASVs) compared to winter (Fig. 4A). We observed greater archaeal variability during summer near the soil-surface (Fig. 4B). In addition, distinct depth-dependent seasonal shifts in the bacterial community composition were noted (Fig. 4C). A detailed description of the general microbial community composition (section 1.1), overall diversity analysis (section 1.2) and seasonal shifts in microbial community composition (section 1.3), are presented in Supplementary Results.

**Figure 4.**
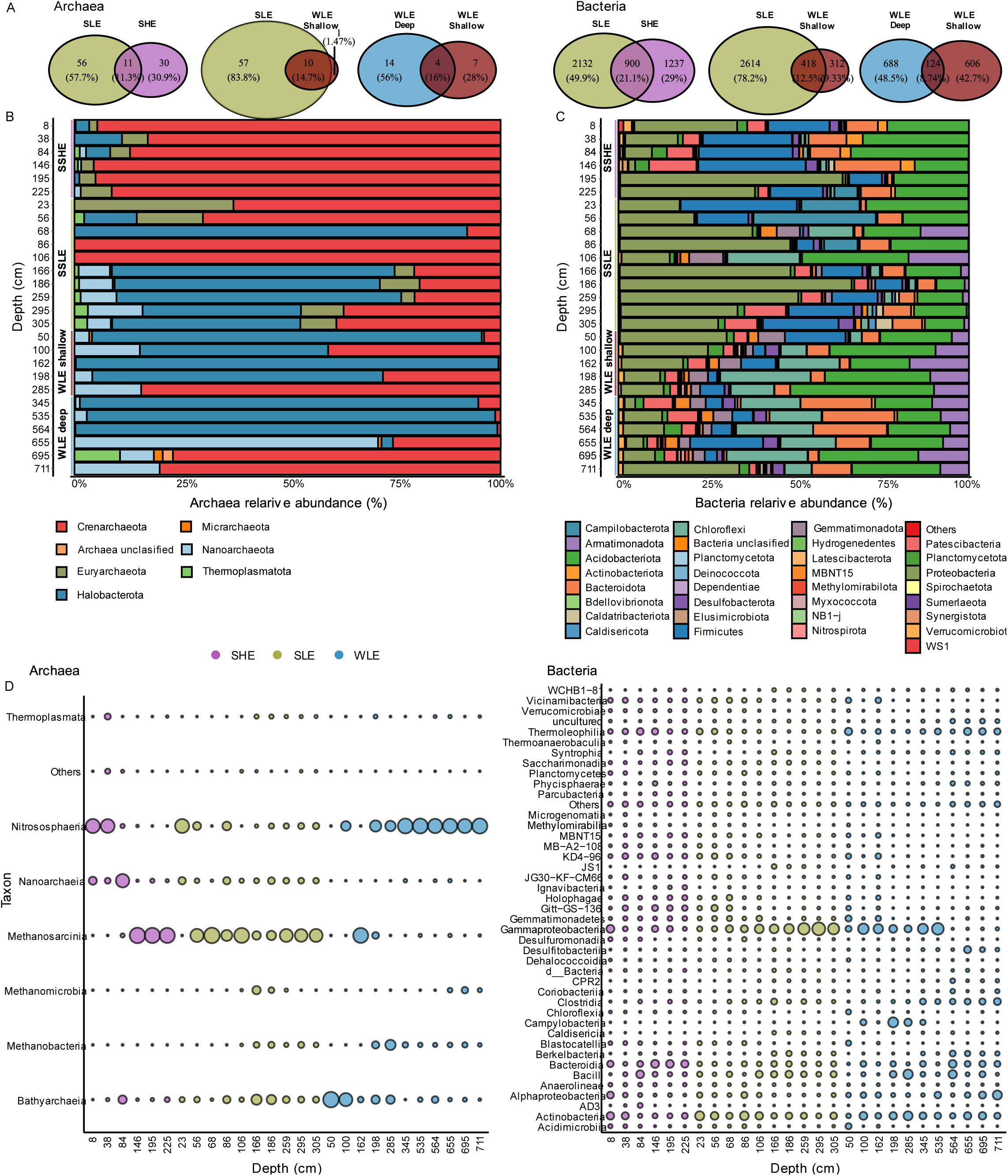
Shifts in microbial community composition at North Star Yedoma boreholes. Analysis of community composition was based on 16S rRNA gene amplicon-based sequencing. Amplicon Sequence Variants (ASVs) were pre-filtered to a retain a minimum frequency of 20 reads, across a minimum of two samples. Three cores were included in the analysis. SME = Summer Mid Elevation (borehole ID: BH1, n=10, olive-green), SHE = Summer High Elevation (BH2, n=6, purple), WME = Winter Mid Elevation (BH6, n=11, blue). When analyzing the full talik, WME samples were further subdivided into shallow (<3 m, n = 5, brown) and deep (>3 m, n = 6, blue) samples (A–C). Summer sampling refers to cores taken on September 15, 2021 and winter on March 18, 2023. Throughout the figure, archaea ASVs are presented at the left panel of each section, while bacteria ASVs at the right. **(A)** Venn diagram visually representing the number of shared and unique ASVs. Three comparisons were made, between: 1. High and mid elevation (SHE vs. SME, at the top 3 m), 2. Seasonal comparison of Summer vs. winter (SME vs. WME, at the top 3 m), and 3. WME Shallow vs. deep samples of the complete talik. **(B+C)** Relative abundance barplot visualization of archaeal **(B)** and bacterial **(C)** ASVs (minimum frequency ≥1%). Samples are ordered on the Y axis according to borehole and depth. **(D)** Bubble plot representing class level relative abundances of archaea (left panel) and bacteria (right panel). Samples were ordered on the X axis according to borehole and depth. Bubble size and color indicate relative abundance and borehole.

The methanogenic community composition in both seasons also exhibited significant depth depended changes, that corresponded to the CH_4_ levels and gene expression profiles. During summer, a diverse methanogenic community was detected mainly between 166-186 cm depths, with PICRUSt2’s K pathway prediction analysis hydrogenotrophic, acetoclastic and methylotrophic methanogenesis. In comparison, winter methanogens were present at deeper depths between 198-285 cm, at much lower relative abundances and comprised of *Methanobacterium* and *Methanosarcina* (Fig. 5A, C and Supplementary Data 5-7). he winter’s methanogenic community was predominantly located at deeper soil layers (655-711 cm) as described in the Axis-2 section.

**Figure 5.**
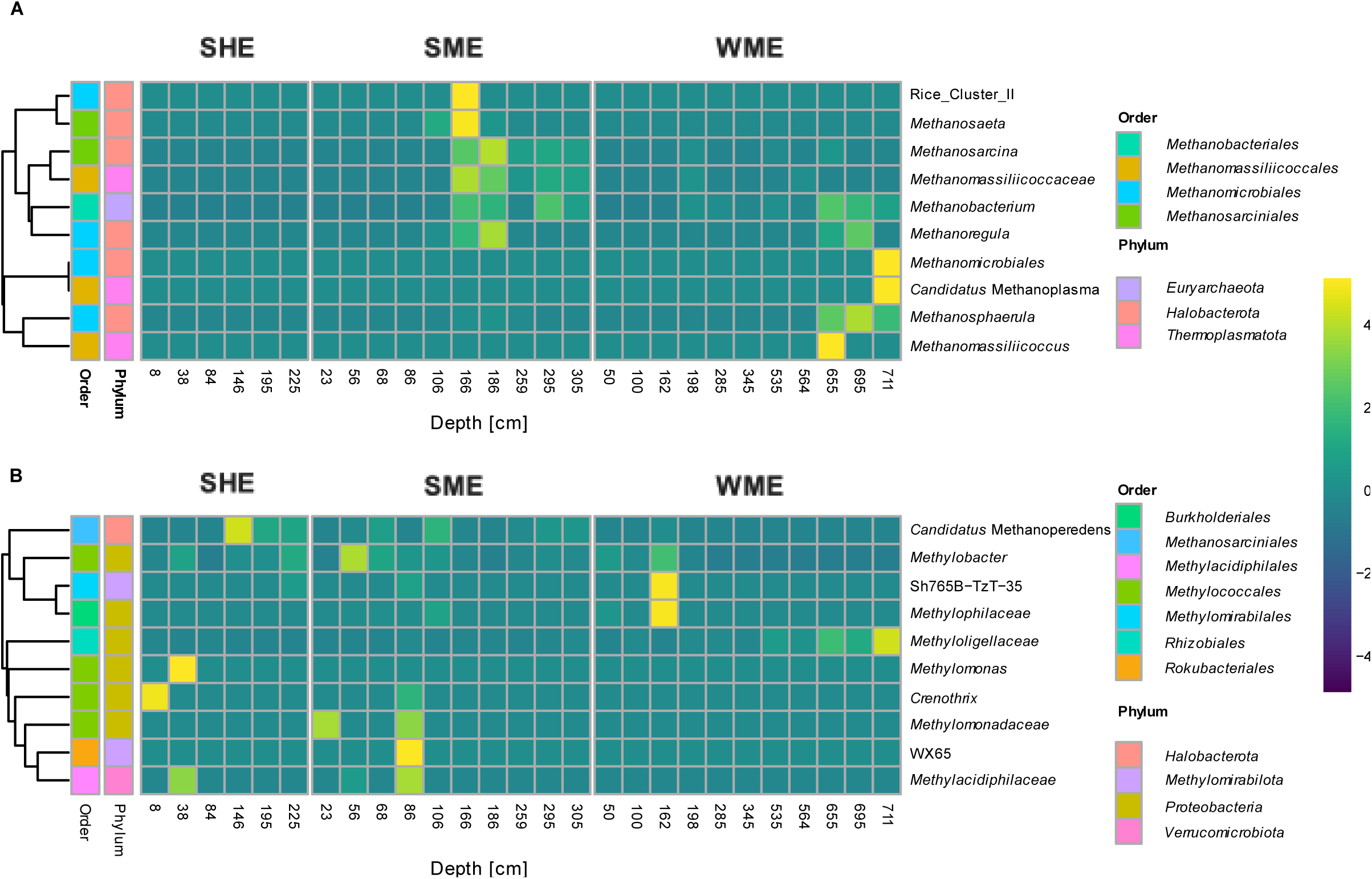
Shifts in the methanogenic and methanotrophic microbial communities at North Star Yedoma boreholes. Analysis of microbial community composition was based on 16S rRNA gene amplicon-based sequencing. Amplicon Sequence Variants (ASVs) were pre-filtered to retain a minimum frequency of 20 reads, across a minimum of two samples. ASVs were collapsed to retain only methanogenic and methanotrophic taxa (rows). Heatmap visualizations present relative abundance of methanogenic **(A)** and methanotrophic **(B)** taxa. For each taxon Order and Phylum are presented to the left of the heatmaps. Samples (columns) were ordered according to depth and faceted according to sampling environment. SME = Summer Mid Elevation (borehole ID: BH1, n=10, olive-green), SHE = Summer High Elevation (BH2, n=6, purple), WME = Winter Mid Elevation (BH6, n=11, blue). The heatmap color scale represents the relative abundance of ASVs within each sample, ranging from low (blue to black) to high (yellow to green), according to the color bar.

During summer, at and above the methanogenic zone, at shallower anaerobic depths, N-AOM coupled to NO_3_^-^ and NO_2_^-^ reduction was exemplified by the high prevalence of *Candidatus* Methanoperedens (ANME2d) (Fig. 5B and Supplementary Data 5). Very high relative abundance was noted at 106 cm (5.5% of all ASVs), but also at 68 cm (5.9%). The high relative abundance at 68 cm is surprising giving the semi aerated conditions. However, given that the oxygen probes were located at specific depths, at thermokarst-mound flanks adjacent to the EC tower in the vicinity of the boreholes, it is likely these did not fully reflect the oxygen concentrations. The high ANME2d relative abundance indicates significant N-AOM in this environment. Contrary, a significant seasonal shift signifying and a reduction in N-AOM was observed during winter, as ANME2d relative abundance plummeted to very low levels. ANME representatives, including *Candidatus* Methanoperedens, have been previously reported in permafrost environments [38, 39, 71]. Coupling AOM to NO_3_^-^ / Fe^3+^ reduction [72], they perform reverse methanogenesis via the mcrA gene and are suggested as potential mitigators of CH_4_ emissions [73, 74]. Yet, at the same time, *Candidatus* Methanoperedens has been related to direct increased N_2_O generation and emissions [75]. We identified a peak in N_2_O at 106 cm, that depleted at 86 cm indicating rapid consumption. We detected at this depth (and beyond) high relative abundance of the genus *pseudomonas* (7.1% at 86 cm). Members of this genus can perform complete aerobic denitrification, and were shown to possess genes catalyzing the reduction of N_2_O to N_2_ [76–78]. *Pseudomonas* can also oxidize CH_4_, and perform AOM with NO_3_^-^ and NO_2_^-^ [79, 80]. The later, together with increased N availability were previously reported following permafrost thaw, culminating in elevated N_2_O emissions [81–83].

Just below the ANME2d maximum peak, we also identify via qPCR the presence of the NC10 phylum, at similar levels at both seasons (Fig. 3). We identified the genus Sh765B-TzT-35 (*Methylomirabilaceae*) in the microbial community analysis, albeit at low relative abundance (Fig. 5B and Supplementary Data 5). Members of this family are known to perform aerobic CH_4_-oxidation (via an intra-aerobic pathway) under anaerobic conditions, coupled to NO_2_^-^ reduction [43]. The NC10 phylum has been previously detected in permafrost peatlands, in mid soil layers where redox gradients and nitrogen availability supported their activity [84]. This highlights their potential role in influencing biogeochemical processes in climate-sensitive systems. Microorganisms like ANME2d and NC10 oxidize CH_4_, coupled to nitrogen species reduction, N_2_O may also be generated through interactions with other members of the microbial consortia [43]. Above these depths, and close to the soil-surface summer methanotrophy was high (68 cm), diminishing by 30-fold in winter-time (Fig. 3 and Table S5), probably due to inhibition related to the near 0 °C temperatures [37]. Indeed, elevated temperatures enhance the expression of both genes [85–87]. As in the upland yedoma domain, similar depth-related patterns in pmoA and mcrA gene expression have been reported in the Arctic and discontinuous permafrost-affected regions [86, 88, 89]. While Higher mcrA levels are typically reported at deeper anaerobic horizons, pmoA expression peaks closer to the soil surface. Shifts in microbial community dynamics (e.g., community composition and spatial distribution) of methanogens, methanotrophs and microorganisms related to the nitrogen cycle, have also been previously reported and associated with changes in environmental conditions such as temperature, pH, water content, soil structure, oxygen level, nutrient availability, carbon and nitrogen content, thaw depth and permafrost collapse [22–24, 29, 34–36, 87–93].

### Seasonal shifts in genes related to denitrification and ammonium oxidation

The observed significant N-AOM at and above the methanogenic zone, predominantly during summer. indicating potential couplings between the CH_4_ and nitrogen cycles. To further examine the potential for N_2_O production, we measured norB absolute gene expression. Similar to the N_2_O concentrations, we identified two peaks at the top 1 m, only in during summer (Figs. 3 and Table S5). These observations raise concerns regarding potential elevations in N_2_O emissions, as previously reported from thermokarst mounds in North Siberian yedoma permafrost [49]. However, unlike NSY with its extended talik formation, North Siberia soils are thawed only several centimeters deep during summer and lack taliks entirely.

Additional nitrogen-cycle genes showed similar absolute expression patterns, with elevated levels closer to the soil-surface in summer (up to 86 cm) (Fig. 3 and Table S5). Thereafter absolute abundance levels dropped sharply. These results indicated processes related to denitrification (narG and nirK genes), anaerobic ammonium (NH_4_^+^) oxidation to N_2_ (Anammox, hzsB) and to NO_2_^-^ (Feammox: 16S of *Acidimicrobiaceae* sp. strain A6). Below these semi-aerated depths, aerobic oxidation of NH_4_^+^ to NO_2_^-^ by bacteria and archaea (amoA gene), peaked between 23-56 cm depths. During winter expression was low for nearly all genes, with slight elevations between 100-162 cm. Elevated expression was noted for nirK and NC10 (Fig. 3 and Table S5), suggesting increased denitrification and methanotrophy coupled to NO_2_^-^ reduction at these anaerobic depths. Pathway prediction analysis, related to the nitrogen cycle also indicated elevated levels near the surface for the summer samples (up to 86 cm depth; Fig. S4B). This included nitrification, denitrification, dissimilatory nitrate reduction and complete nitrification, comammox. Contrary, winter nitrogen related pathways were predicted for the most-part at very low levels, with slight elevations at deeper sediment depths (∼162 cm, Fig. S4B and Supplementary Data 6-7). Winter denitrification was predicted at relatively high levels as during summer.

Our data indicates significant seasonal shifts in microbial community dynamics and functional, related to both the CH_4_ and nitrogen cycles. While during winter we observed higher CH_4_ emissions, due to soil-surface freezing that neared the freezing point. During summer, we observed N-AOM, together with elevated N_2_O generation. Continued warming is projected to drive widespread permafrost thaw and talik formation across the yedoma domain, releasing vast pools of old carbon and nitrogen [5, 93]. These can become accessible to microbial mineralization and stimulation of GHG emissions, including N_2_O [11, 35, 93]. The latter has been related to the activity of denitrifying microorganisms at the top surface layers [49, 94, 95]. Furthermore, winter warming may lead to deepening of the active layer and talik formation, increased water permeability, re-vegetation growth, and soil mixing after permafrost collapse [31, 64, 93, 96]. To date, most studies represent continuous-permafrost or non-yedoma environments. Augmented permafrost thaw can alter soil temperature, water retention, drainage and soil-moisture [97], in a way that is difficult to predict on a large-scale basis. Our NSY upland study site is characterized by relatively dry soil conditions, with GWC ranging between 18%-32% (Table S1 and Walter Antony et al., 2024 [37]). Previous reports indicate intermediate moisture content promotes N_2_O emissions from yedoma and other permafrost affected soils [50, 67]. Continued global warming may lead to elevated GWC in upland yedoma areas of continuous-permafrost, potentially facilitating the establishment of denitrifying and N_2_O producing microbial communities augmenting GHG emissions [49]. With recent data suggesting that nitrogen mineralization and turnover rates in permafrost-affected active layers are comparable to those in temperate and tropical soils [10], our findings underscore the importance of microbial and biogeochemical dynamics related to widespread permafrost degradation and talik formation across the yedoma domain. These seasonal shifts highlight the sensitivity of permafrost systems to climatic changes and their potential augmentation of global GHG emissions.

### Axis 2 – Depth profile of the full talik down to the top of permafrost

The yedoma domain, though small in area, contains a large share of the northern permafrost soil C_org_ and N_org_ pools extending tens of meters deep [1, 5]. Heat transport within the thawing yedoma permafrost, further accelerates talik formation, facilitating microbially mediated organic matter mineralization [37]. Analyzing the depth profile of the full talik down to the top of permafrost at NSY (Axis 2), provides an opportunity to study CH4 and nitrogen related microbial processes linked to GHG emissions, particularly in the context of global warming and talik formation.

The only full-talik-profile core we obtained was in winter. Our PCA analysis of the winter’s (WME) full talik profile, showed an overlap between shallow (≤3 m) and deeper (>3 m) soil layers, with great variability in most physio-chemical parameters (Fig. 2B and Table S4). In addition to the first CH_4_ peak observed at the shallower depths (285 cm), a second peak was measured at deeper layers (535-655 cm). This was accompanied by elevations in mcrA expression, with two peaks at 285 cm and at 711 cm. This offset at the deeper layers, may be related to upward gas migration [37]. The absolute expression levels of all other genes related to aerobic methanotrophy and the nitrogen cycle were low, across all deeper depths (Fig. 3 and Table S5). In addition, we did not identify any N_2_O production in the winter core.

Although shallow and deep samples presented a similar proportion of unique bacterial ASVs (48.5%-42.7%), that of archaea was higher in the shallower samples (56% vs. 28%), yet for the latter, species richness was low (Fig. 4A). The dominated archaea phylum in both environments was *Crenarchaeota* (52% and 90%, respectively; Fig. 4B, D and Supplementary Data 3), at almost all depths. While the methanogenic community in the shallower depths mainly consisted of *Methanobacterium* and *Methanosarcina* at low relative abundance, in deeper depths a more diverse community was noted (Fig. 5A and Supplementary Data 5). Pathway prediction analysis supported mainly acetoclastic methanogenesis at both depths (Fig. S4A and Supplementary Data 6-7). The methanotrophic community throughout the borehole was less diverse (Figs. 5B and Supplementary Data 5, see below for elaboration) and aerobic methanotrophy pathway prediction was very low (Fig. S4A). Significant changes in bacterial community composition were also observed (Fig. 4C, Supplementary Data 4). A more detailed description of Axis-2 microbial community composition is presented in Supplementary Results 1.4.

Our results indicate significant differences in both archaea and bacterial community dynamics, between shallow and deeper depths of the winter full talik profile. The diminished CH_4_ and nitrogen-related processes at the deeper talik horizons, compared to the soil surface, may be related to lower temperatures and functional constraints, leading to decreased N-AOM, higher CH_4_ emissions and lower N_2_O production. Microbial communities at deeper, more ancient yedoma permafrost deposits may be shaped by long-term cryogenic conditions, exhibiting specialized survival strategies for persistence in the frozen, isolated, resource-limited environment [23]. Additional constraints on carbon and nitrogen cycling in these ecosystems have been attributed to missing microbial functions [93]. While freshly thawed wet yedoma sediments exhibit low N_2_O emissions due to such limitations on key microbial functional groups, following long-term thaw, significant shifts can alter microbial community composition and function, resulting in higher N_2_O emissions [49].

### Axis 3 – Elevation and aeration driven alterations in microbial dynamics in the CH_4_ and nitrogen cycles

During summer, we analyzed two boreholes, characterized by high (SHE) and mid (SME) elevation, corresponding to deep vs. shallow aeration conditions. In contrast to the winter borehole which extended to 711 cm depth, these boreholes extended only down to 225 cm at SHE and 305 cm at SME. Among our summer samples, we observed distinct groupings with partial overlap in the PCA analysis between samples. In the top 1.5 m samples group together, characterized by low CH_4_ and elevated CO_2_ concentrations (Fig. 2A-B and Table S1,4). Beyond these depths, SME samples showed higher CH_4_ and lower CO_2_ concentrations. methane concentrations in the SHE core were low throughout, peaking at 225 cm (0.06 mM). Corresponding patterns were noted in the mcrA and pmoA gene absolute expression. Lower mcrA levels were measured for the SHE samples, across most depths (Fig. 3 and Table S5). An increase in mcrA expression, restricted to 225 cm depth was noted, supporting the existence of a deeper aerobic zone. In both environments, aerobic and anaerobic methanotrophy was significant. pmoA peaked near the soil-surface (8 cm), while CH_4_ was absent at top-soil aerated layers, as in SME samples. One N_2_O peak was noted in the SHE samples (2.7 µM at 91 cm), that was 2.6 times lower compared to the SME samples. Yet, norB expression in SHE was below detection limit at all depths. These differences may be related to increased N availability at lower altitudes [98, 99], or to other contributing factors indirectly related to elevation, including temperature, soil and water chemistry, oxygen-availability and microbial activity [100, 101]. Nitrogen-cycle related genes of the SHE samples, followed a similar expression pattern to that of SME, peaking closer to the soil-surface (Fig. 3 and Table S5), followed by sharp declines.

SME samples contained twice as many unique bacterial and archaeal ASVs (Fig. 4A), indicating higher diversity. Depth dependent differences were detected in both boreholes in bacteria and archaea phyla. A detailed description of axis-3 microbial community composition is presented at Supplementary Results 1.5.

In agreement with the CH_4_ profile and supporting more aerated conditions, methanogens were absent from the SHE borehole, up to 225 cm (Fig. 5A and Supplementary Data 5). These, together with the relatively low CH_4_ concentrations and observed positive emissions in chamber fluxes [37], suggest that a methanogenic community is likely present at deeper layers. Despite low methane concentrations, pmoA absolute expression (Fig. 3) and aerobic methanotrophs were abundant closer to the soil-surface (Fig. 5B and Supplementary Data 5). pmoA and amoA are evolutionary related [102], and portray several structural and functional similarities [103]. In nitrogen-rich environments, this may enable aerobic methanotrophs with denitrification capabilities to oxidize ammonium, and perform incomplete denitrification [104]. Permafrost thaw and subsequent increased nitrogen availability, may contribute to elevated N_2_O emissions [49, 105].

As with the SME core, we identified *Candidatus* Methanoperedens at very high levels in the SHE core, at 146 cm (9.2% of all ASVs) and 195 cm (5.9%) depths (Fig. 5B and Supplementary Data 5; discussed below). No methanogens were identified in our analysis of SHE samples. Yet, methanogenesis was exemplified by qPCR of the mcrA gene and PICRUSt2’s pathway prediction analysis (Fig. 3 and S4A and Supplementary Data 6-7). It is probable that the results reflect the reverse methanogenesis performed by ANME2d.

### Methane dynamics in upland yedoma

In this study we delineate the spatio-temporal dynamics of methanogenic and methanotrophic communities in upland yedoma deposits affected by permafrost thaw and talik formation. Our findings highlight the predominance of methanogenesis and couplings between methanotrophy and the nitrogen cycle, and their potential implications for GHG emissions. During summer, under mid elevation and a shallower aerobic depth zone, CH_4_ production was higher as methanogens were more prevalent at anaerobic depths closer to the soil-surface. Summer methanotrophy was augmented by aerobic and anaerobic methanotrophs (i.e., ANME2d, NC10 and denitrifiers performing N-AOM). We observed two distinct N_2_O peaks within the top 1 m. The deeper of these, coincided with a high abundance of ANME-2d archaea, associating it to CH_4_ oxidation. The second N_2_O peak, located closer to the soil surface, was of a similar magnitude. Given that N_2_O is approximately nine times more potent as GHG a than CH_4_, the contribution of this surface-associated peak to the overall GHG emissions effect is expected to be significant. During winter, near freeze temperatures at the soil-surface, limited microbial activity, and methanotrophy. Anaerobic N-AOM was reduced, and ANME2d were largely absent, possibly due to near freeze temperatures and reduced nitrogen availability during winter. We did not observe N_2_O generation in winter, a time whenCH_4_ emissions were high. In summer, we found evidence for N_2_O generation up to ∼1 m depth, coupled to denitrification and N-AOM by *Methanoperedens*. These were absent during winter, probably due to decreased temperatures and additional related processes such as nitrogen availability and other environmental factors.

These mechanisms demonstrate the dynamic interplay between aerobic and anaerobic processes of the CH_4_ and nitrogen cycles. The suggested mechanisms are illustrated in Fig. 6, based on the chemical, qPCR and 16S amplicon-based sequencing analyses.

**Figure 6:**
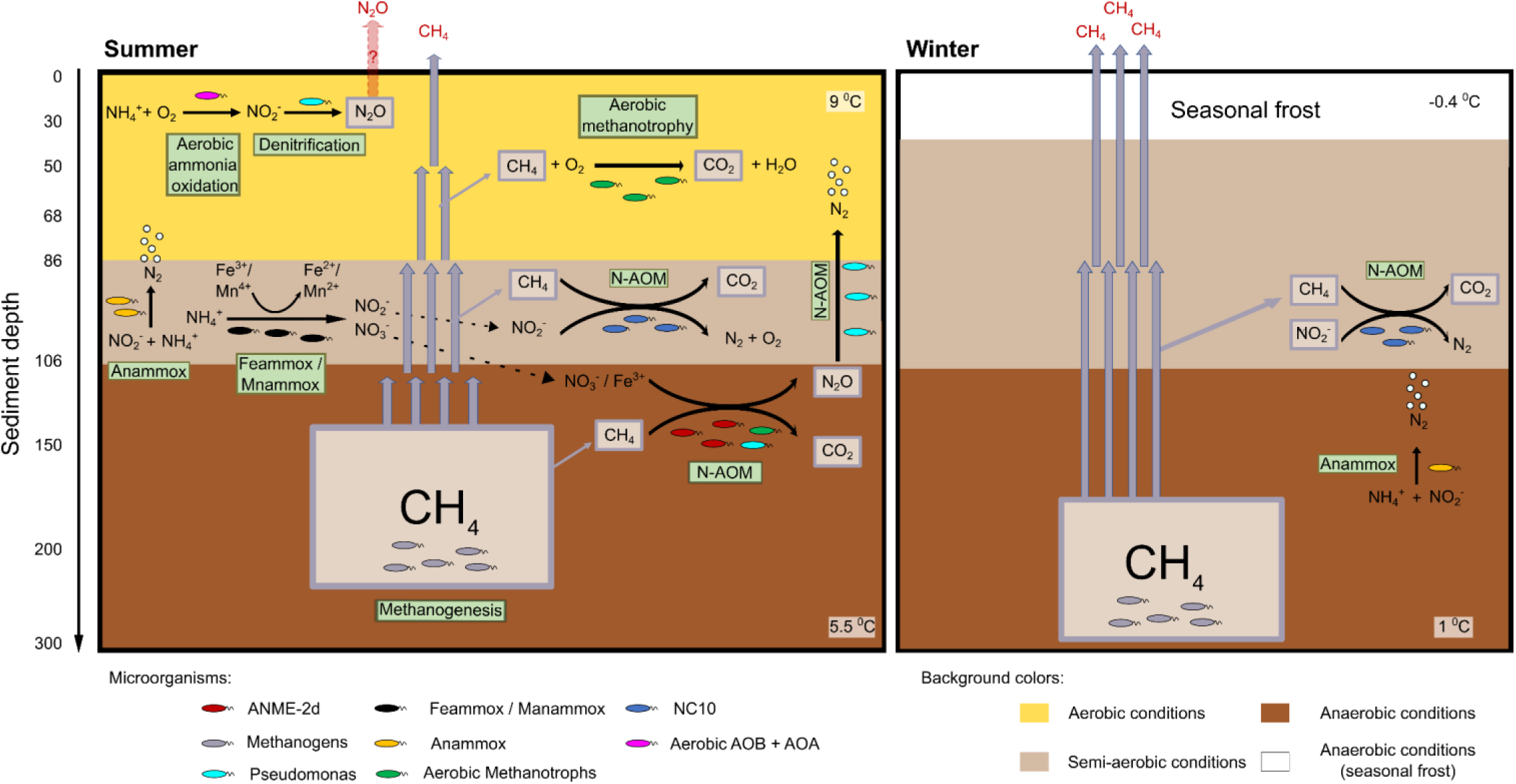
Methane dynamics in upland yedoma, suggested mechanisms. During summer (left panel), methanogenesis occurs in deeper anaerobic talik layers, with CH_4_ oxidized by both anaerobic (at deeper) and aerobic (at shallower layer) methanotrophy, by ANME2d, NC10, and denitrifiers performing N-AOM. methane oxidation is coupled to the nitrogen cycle, with N-AOM linking CH_4_ and nitrogen transformations. N_2_O is present up to 1 m depth, with its distribution influenced by microbial activity. In winter, near-freezing temperatures at the soil surface limit microbial activity and methanotrophy. The diminished CH_4_ oxidation and nitrogen-related processes at deeper talik horizons, compared to the surface, may be related to lower temperatures, functional constraints, or nitrogen availability, leading to decreased N-AOM, higher CH_4_ emissions, and lower N_2_O production. This figure illustrates the seasonal dynamics of CH_4_ and nitrogen cycling, highlighting the interplay between methanogenesis, methanotrophy, and nitrogen transformations in thawing permafrost. The suggested mechanisms are based on chemical, qPCR, and 16S amplicon-based sequencing analyses from this study. The background colors represent different soil oxygenation conditions. Yellow indicates aerobic conditions. The light-brown areas correspond to semi-aerated conditions, and the dark-brown areas represent fully anaerobic conditions, where oxygen is absent, and strictly anaerobic processes dominate. The white area illustrates winter seasonal frost, with anaerobic conditions near the soil surface (top 15 cm). In-depth mechanistic analysis and description of oxygen mobilization between the top soil layers is presented in our previous paper [37].

Microbial communities in permafrost soils are important players in regulation of carbon and nitrogen cycling, particularly as thaw releases the frozen organic matter. Thawing of yedoma permafrost may liberate significant nitrogen stocks, that are predicted to contribute to denitrification and N-AOM, potentially elevating N_2_O emissions [5, 11, 49, 50]. GHGs are produced by microbial activity, that is strongly influenced by the microbial composition and its functional capacities [67, 93]. The microbial processes in yedoma soils are often limited by the functional constraints imposed by prolonged freezing over millennia. Thawing may alleviate some of these constraints by introducing functionally diverse microbial communities, facilitating carbon and nitrogen cycling [11, 93] and augment aerobic and anaerobic methanotrophy at deeper layers, including N-AOM. These are influenced by factors like moisture, oxygen availability, and temperature [49, 106]. Permafrost carbon and nitrogen feedbacks/interactions/couplings in yedoma soils, and the role of nitrogen cycling, particularly processes like N-AOM, remain poorly understood [49, 50].

### Concluding remarks

The interplay between permafrost thaw, alleviation of carbon and nitrogen stocks, microbial activity and elevated GHG emissions, accentuate the significance of yedoma permafrost soils. Yet, most studies conducted on yedoma permafrost, focused on the uppermost layers of uplands in the colder, continuous permafrost zone, or on thermokarst lakes. Our data, together with previous flux compilations, show that during both summer and winter, upland yedoma is a surprisingly high source for CH_4_ and potential for N_2_O emissions. Our soil analyses suggest that emissions of N_2_O, unlike CH_4_, may be higher in summer than in winter. Unsaturated-thawed yedoma permafrost harbors a diverse microbial community, with distinct spatio-temporal shifts in composition and function. This is evident in the methanogenic and methanotrophic community dynamics, as well as N-AOM processes. These findings are particularly important under continued global warming, which is expected to cause widespread talik formation in the colder yedoma region of North Siberia by the end of this century and warmer upland yedoma soils everywhere.

## Supporting information

Description of Additional Supplementary Files

## Acknowledgments

Site history and access to the North Star Yedoma field site were provided by Roger and Melinda Evens and Raymond and Stephanie Nadon. Peter Anthony, Nicholas Hasson, and Colin Edgar assisted with fieldwork.

## Author contributions

O.B., K.M.W.A., and O.S. Conceived the study. O.B. conducted the microbial analyses on soil cores, performed the statistics, and wrote the paper. K.M.W.A. led the field work and core collection. O.S., and E.E.R. Performed the biogeochemical profiles. All authors, commented on the analysis, interpretation, and presentation of the data, and were involved in the writing.

## Data availability

All the data used to generate the figures and analyses in this study are included in this published article, supplementary information and supplementary data files. The datasets and scripts that were used for the analyses presented in the manuscript will be applauded upon acceptance to the GitHub repository and will be made available. Raw metagenomic sequences reads generated in this study have been deposited in in the European Nucleotide Archive Database (https://www.ebi.ac.uk/ena/browser/home) as project accession number PRJEB79303, Secondary Accession ERP163479. Additional data are available under the Supplementary Information and Supplementary Data sections.

## Code availability

The datasets and scripts that were used for the analyses presented in the manuscript will be applauded upon publication to the GitHub repository and will be made available.

