## Supplementry information for "Complex nitrogen redox couplings control methane emissions from Arctic upland yedoma taliks"


**The following are included:**

**Supplementary Figs 1-4**

**Supplementary Tables 1-11**

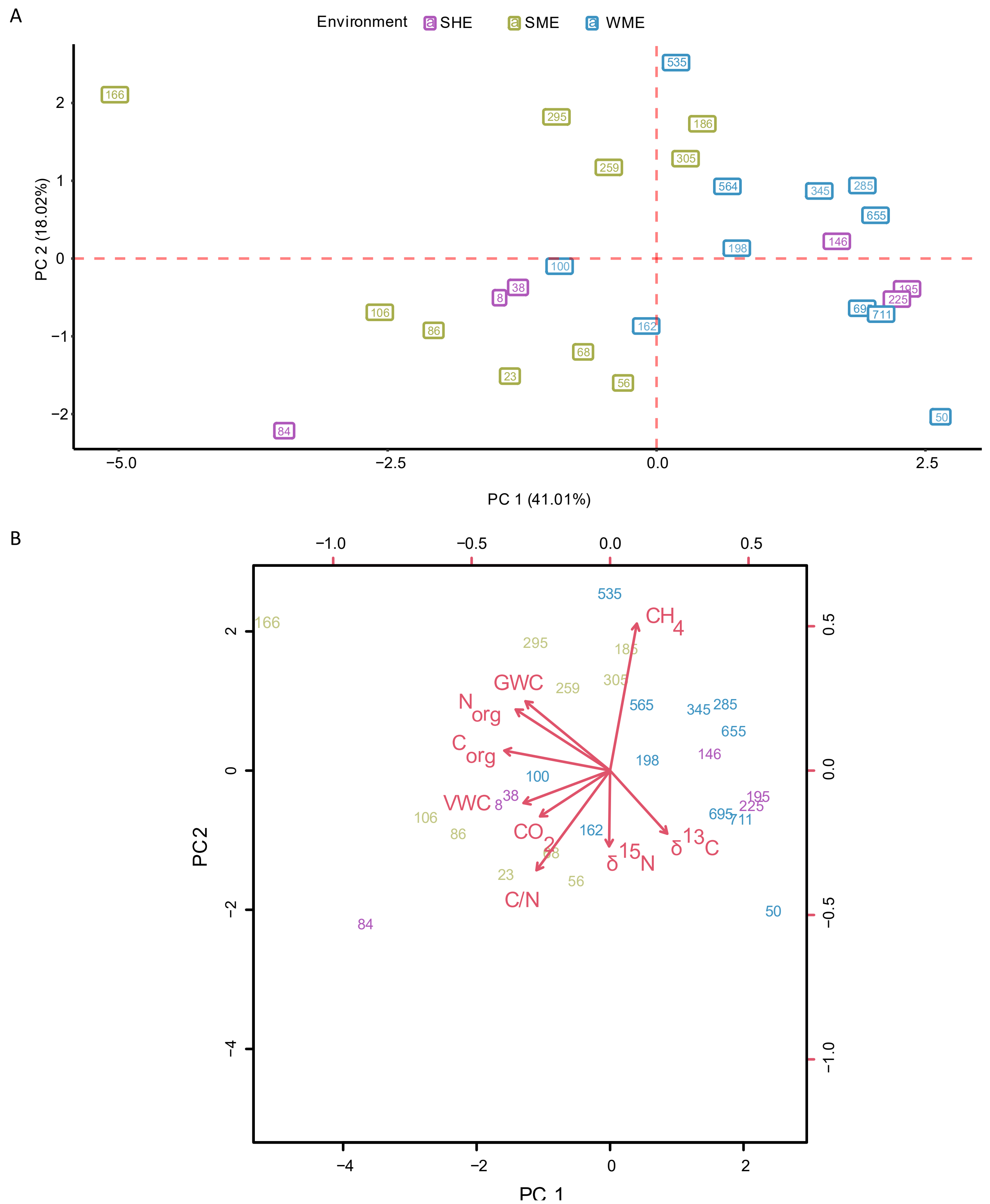


**Figure S1: Initial PCA analysis of the environmental parameters at the NSY study site.** (A) The initial PCA included 9 environmental parameters (Table S1): Volumetric Water Content (VWC)

Gravimetric Water Content (GWC), $CH_4$ and $CO_2$ concentrations, organic Carbon ($C_{org}$) (%), organic Nitrogen $N_{org}$ (%), $\delta^{15}N$ (‰), $\delta^{13}C$ (‰). To maintain the three main axes of comparison, we have denoted the sampled cores by season and elevation, as follows: SME = Summer Mid Elevation (borehole ID: BH1, n=10, olive-green), SHE = Summer High Elevation (BH2, n=6, purple), WME = Winter Mid Elevation (BH6, n=11, blue). Summer sampling refers to cores taken on the September 15, 2021 and winter on March 18, 2023. **(B)** Loading scores, indicating the importance of tested environmental variables related to PC 1 and PC 2. Based on collinearity and the loading scores (Tables S2-S3) of this initial PCA analysis, a final PCA was performed (Fig. 2).

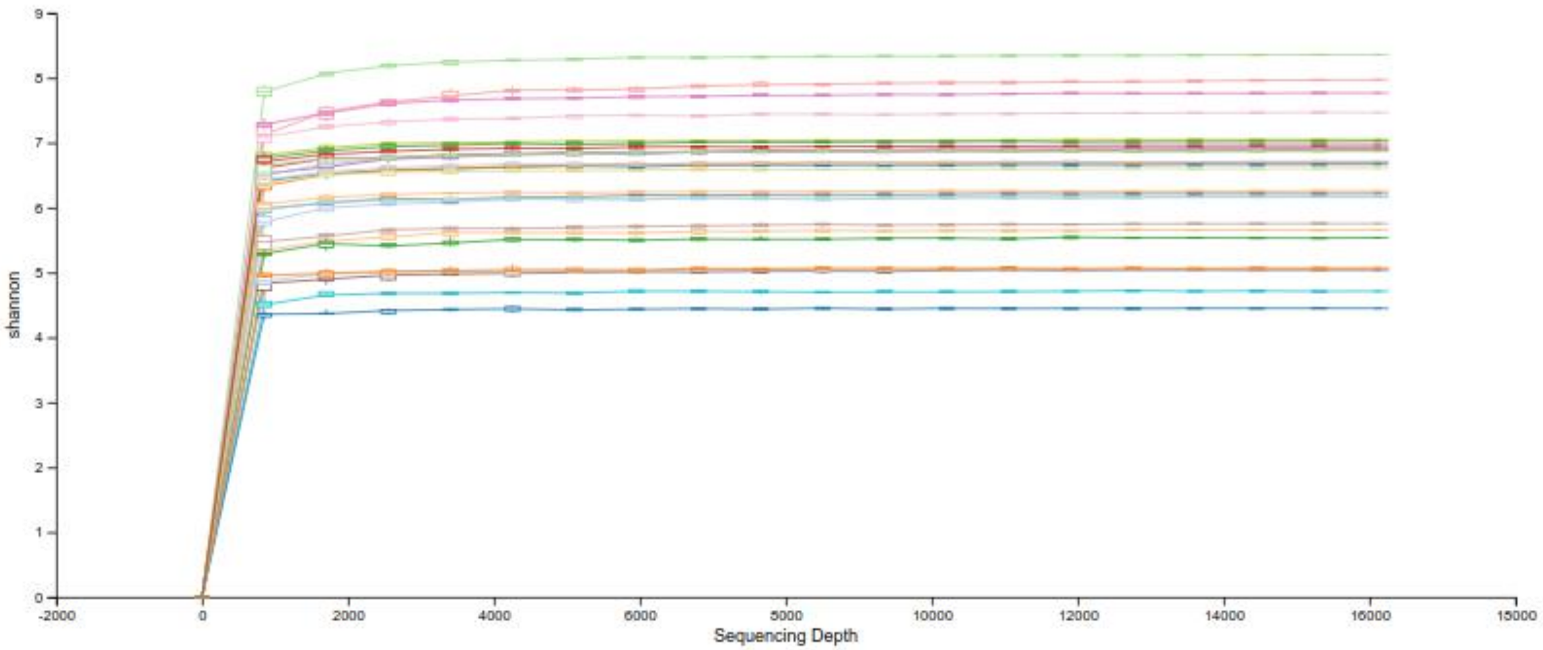


**Figure S2. Rarefaction curves for 16S rRNA gene amplicon-based sequencing.** Amplicon Sequence Variants (ASVs) were pre-filtered to a minimum frequency of 20 reads across all samples, in a minimum of 2 samples. Rarefaction curves were generated using the q2-alpha-rarefaction function of the q2-diversity plugin. The colors represent the different samples.

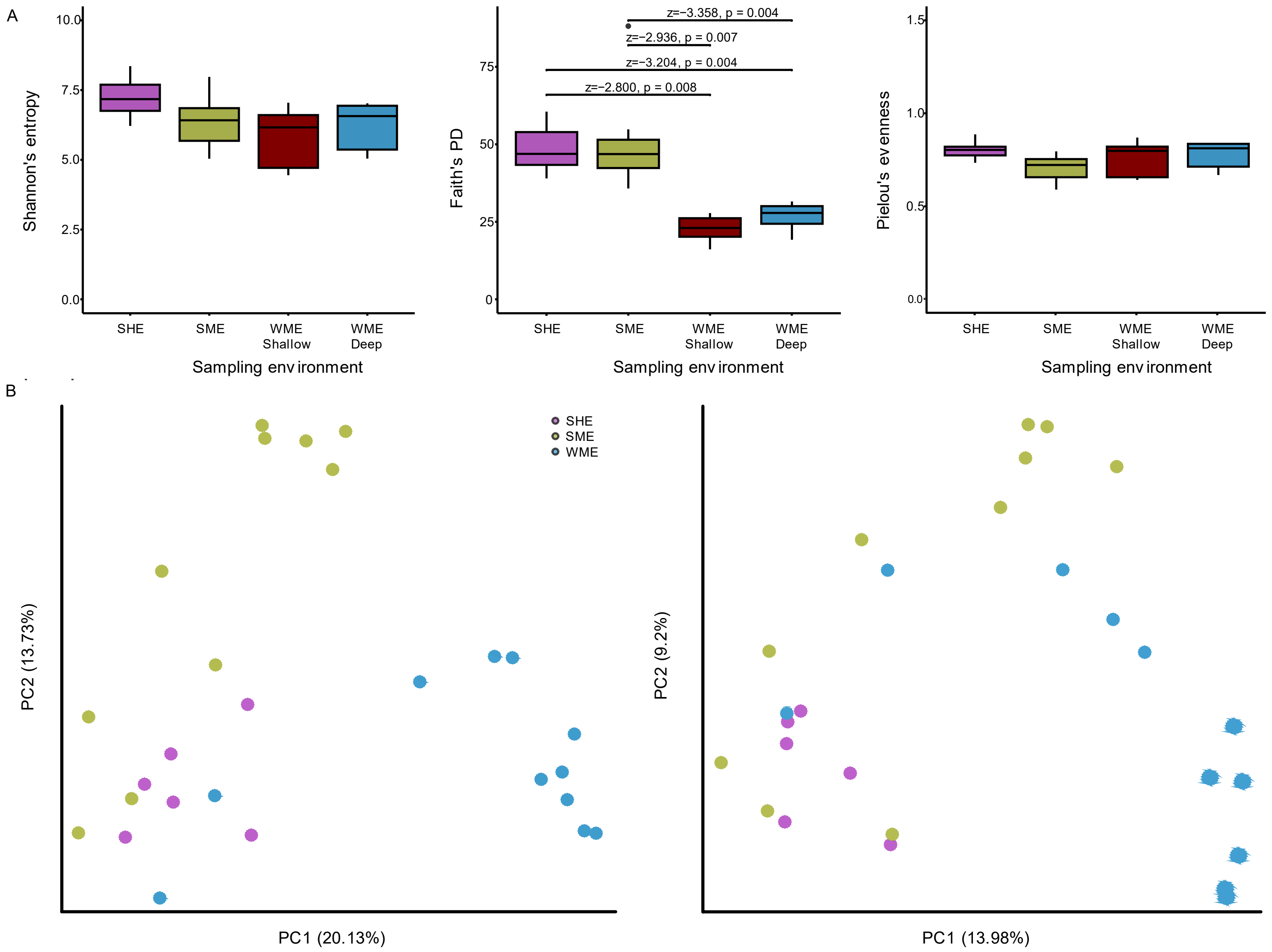


**Figure S3. Microbial diversity North Star Yedoma (NSY).** Based on 16S rRNA gene amplicon-based sequencing. Amplicon Sequence Variants (ASVs) were pre-filtered to a minimum frequency of 20 reads across all samples, in a minimum of 2 samples. **(A)** Alpha diversity analysis: Shannon's entropy (left panel), Faith's PD (midle panel) and Pielou's evenness (right panel) were included in the analysis. Kruskal-Wallis test results and post hoc via Wilcox tests are presented in Supplementary Tables S6-S7. **(B)** Beta diversity was assessed via Jaccard (left panel) and Bray–Curtis (right panel). Statistical significance was tested via PERMANOVA and PERMDISP analyses (Tables S8-S9). To maintain the three main axes of comparison, we have denoted the sampled cores by season and elevation, as follows: SME = Summer Mid Elevation (borehole ID: BH1, n=10, olive-green), SHE = Summer High Elevation (BH2, n=6, purple), WME = Winter Mid Elevation (BH6, n=11). For comparison of alpha diversity full talik WME samples were subdivided to shallow (n=5, brown)

and deep (n=6, blue) samples. Summer sampling refers to cores taken on the September 15, 2021 and winter on March 18, 2023.

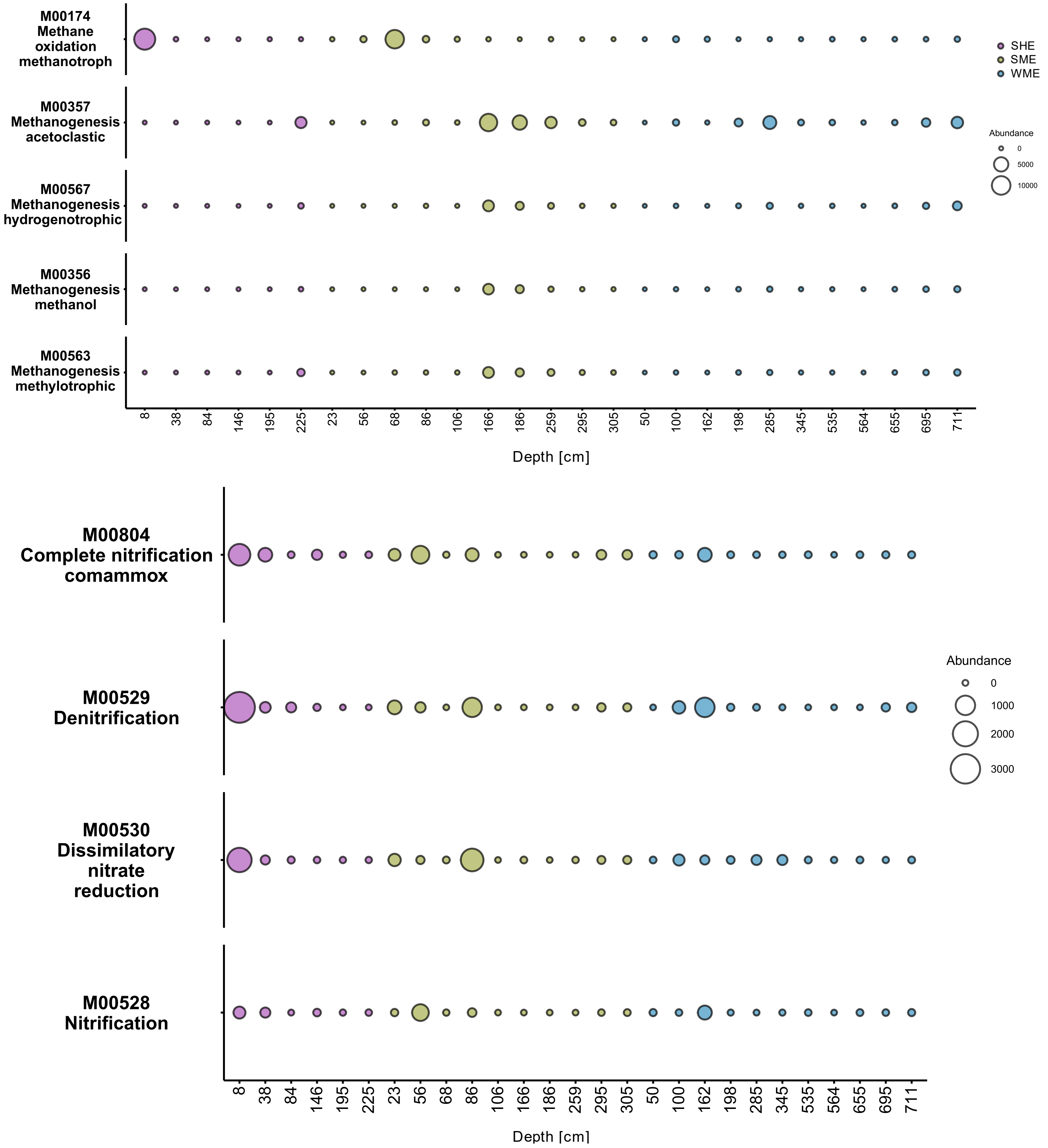


**Figure S4. Figure S3. The abundance of $CH_4$ and nitrogen cycles pathway prediction analysis.** Based on 16S rRNA gene amplicon-based sequencing. Amplicon Sequence Variants (ASVs) were pre-filtered to a minimum frequency of 20 reads across all samples, in a minimum of 2 samples. Raw data was analyzed with the PICRUSt2 package, corrected for qPCR resulats (see material and methods section for a detailed description). (A) Methane pathway prediction analysis and (B) Nitrogen pathway prediction analysis. To maintain the three main axes of comparison, we have denoted the sampled cores by season and elevation, as follows: SME = Summer Mid Elevation (borehole ID: BH1, n=10, olive-green), SHE = Summer High Elevation (BH2, n=6, purple), WME = Winter Mid Elevation (BH6, n=11, blue). Summer sampling refers to cores taken on the September 15, 2021 and winter on March 18, 2023.

**Table S1:** Physicochemical characteristics of North Star Yedoma (NSY) samples, sent for microbiological samples. Included variables were: variables volumetric water content (VWC), gravimetric water content (GWC), $CH_4$ and $CO_2$ concentrations, $C_{org}$ (%), $N_{org}$ (%), $C_{org}/N_{org}$ ration, $\delta^{15}N$ (‰), $\delta^{13}C$ (‰). In addition, for each sample the following parameters are presented: Sampling depth, season, stratification period. Samples were characterized based on sampling environments: SHE = Summer Shallow High Elevation (n=6), SLE = Summer Shallow Low Elevation (n=10), WLE = Winter Low Elevation (n=11). The WLE samples were comprised of shallow (<3 m, n=5) and deep (<3 m, n=6) samples. For elaboration, see materials and methods.

| Sample ID | Borehole | Environment | season | Depth (cm) | VWC (%) | $CH_4$ (mM) | $CO_2$ (mM) | GWC (%) | $N_{org}$ (%) | $C_{org}$ (%) | $C_{org}$ / $N_{org}$ (ratio) | $\delta^{15}N$ (‰) | $\delta^{13}C$ (‰) |
|---|---|---|---|---|---|---|---|---|---|---|---|---|---|
| BH2_8 | BH2 | SHE | summer | 8 | 31 | 0 | 5.2 | 25 | 0.16 | 2.13 | 13.6 | 1.7 | 26.3 |
| BH2_38 | BH2 | SHE | summer | 38 | 26.3 | 0 | 7.2 | 23 | 0.22 | 2.62 | 12 | 3.3 | 26.4 |
| BH2_84 | BH2 | SHE | summer | 84 | 37.2 | 0.019 | 12.1 | 26 | 0.16 | 3.32 | 20.2 | 4.5 | 26.8 |
| BH2_146 | BH2 | SHE | summer | 146 | 24.8 | 0.004 | 5.8 | 20 | 0.08 | 0.7 | 8.9 | 0.7 | 25.3 |
| BH2_195 | BH2 | SHE | summer | 195 | 26 | 0.003 | 0.012 | 21 | 0.06 | 0.55 | 9.2 | 4.2 | 25.3 |
| BH2_225 | BH2 | SHE | summer | 225 | 28.5 | 0.061 | 0.085 | 23 | 0.03 | 0.31 | 9.6 | 4.2 | 24.9 |
| BH1_23 | BH1 | SLE | summer | 23 | 38.4 | 0 | 4.1 | 25 | 0.09 | 1.37 | 15.5 | 1.7 | 26.4 |
| BH1_56 | BH1 | SLE | summer | 56 | 32.4 | 0 | 4.9 | 23 | 0.07 | 1.06 | 14.2 | 3.2 | 26 |
| BH1_68 | BH1 | SLE | summer | 68 | 32.9 | 0 | 5.8 | 24 | 0.1 | 1.32 | 13 | 3.1 | 26.8 |
| BH1_86 | BH1 | SLE | summer | 86 | 41.5 | 0 | 4.8 | 32 | 0.11 | 1.45 | 13.5 | 3.1 | 27 |
| BH1_105 | BH1 | SLE | summer | 106 | 35.8 | 0.043 | 7.6 | 29 | 0.17 | 2.5 | 14.3 | 2.6 | 26.7 |
| BH1_166 | BH1 | SLE | summer | 166 | 35 | 0.274 | 5.2 | 32 | 0.31 | 4.03 | 12.9 | 2.7 | 0 |
| BH1_185 | BH1 | SLE | summer | 186 | 27.5 | 1.967 | 4.8 | 26 | 0.09 | 1.01 | 11.5 | 0.4 | 25.6 |
| BH1_259 | BH1 | SLE | summer | 259 | 27.2 | 1.051 | 1.1 | 26 | 0.18 | 2.1 | 11.5 | 2.6 | 25.6 |
| BH1_295 | BH1 | SLE | summer | 295 | 27.6 | 1.572 | 3 | 27 | 0.21 | 2.33 | 11 | 1.8 | 25.9 |
| BH1_305 | BH1 | SLE | summer | 305 | 28.9 | 1.439 | 2.6 | 28 | 0.07 | 0.95 | 12.6 | 0.5 | 26 |
| BH6_50 | BH6 | WLE | winter | 50 | 22 | 0.008 | 0.8 | 15 | 0.03 | 0.51 | 15.06 | 2.98 | 26.17 |
| BH6_100 | BH6 | WLE | winter | 100 | 28 | 0.059 | 0 | 27 | 0.19 | 2.45 | 12.89 | 4.5 | 27.08 |
| BH6_162 | BH6 | WLE | winter | 162 | 30 | 0.047 | 7.7 | 24 | 0.09 | 1.05 | 11.61 | 2.98 | 25.68 |
| BH6_198 | BH6 | WLE | winter | 198 | 25 | 0.474 | 0 | 28 | 0.09 | 1.1 | 12.5 | 4.25 | 25.99 |

| | | | | | | | | | | | | | |
|---|---|---|---|---|---|---|---|---|---|---|---|---|---|
| BH6_285 | BH6 | WLE | winter | 285 | 28 | 2.279 | 0 | 25 | 0.06 | 0.57 | 9.88 | 7.24 | 25.06 |
| BH6_345 | BH6 | WLE | winter | 345 | 29 | 1.539 | 2.7 | 21 | 0.06 | 0.63 | 10.48 | 1.28 | 24.66 |
| BH6_535 | BH6 | WLE | winter | 535 | 30 | 3.017 | 0 | 27 | 0.16 | 1.63 | 10.46 | 1.9 | 25.46 |
| BH6_564 | BH6 | WLE | winter | 564 | 31 | 1.465 | 3.9 | 24 | 0.08 | 0.86 | 10.31 | 1.75 | 25.13 |
| BH6_655 | BH6 | WLE | winter | 655 | 26 | 1.194 | 0 | 21 | 0.07 | 0.76 | 10.62 | 3.44 | 24.98 |
| BH6_695 | BH6 | WLE | winter | 695 | 25 | 0.4 | 3.9 | 18 | 0.06 | 0.66 | 10.73 | 2.95 | 25.25 |
| BH6_711 | BH6 | WLE | winter | 711 | 26 | 0.044 | 0 | 18 | 0.07 | 0.77 | 11.13 | 2.82 | 25.07 |

**Table S2:** Correlations between the physicochemical parameters comprising the initial PCA. Correlations $\geq$ 0.7, with p-values < 0.05 are marked bold with underline. Included variables were: variables volumetric water content (VWC), gravimetric water content (GWC), $CH_4$, $CO_2$, $C_{org}$, $N_{org}$, $C_{org}/N_{org}$ ration, $\delta^{15}N$, $\delta^{13}C$.

| Variable 1 | Variable 2 | cor | p-value |
|---|---|---|---|
| $N_{org}$ | $C_{org}$ | **<u>0.956</u>** | **<u>0.000</u>** |
| C/N | $\delta^{13}C$ | **<u>0.743</u>** | **<u>0.000</u>** |
| GWC | $C_{org}$ | 0.687 | 0.000 |
| GWC | $N_{org}$ | 0.669 | 0.000 |
| $C_{org}$ | C/N | 0.573 | 0.002 |
| $CH_4$ | $\delta^{13}C$ | -0.565 | 0.002 |
| $C_{org}$ | $\delta^{13}C$ | 0.548 | 0.003 |
| VWC | GWC | 0.544 | 0.003 |
| $CH_4$ | C/N | -0.532 | 0.004 |
| VWC | $C_{org}$ | 0.514 | 0.006 |
| VWC | C/N | 0.508 | 0.007 |
| $CH_4$ | $CO_2$ | -0.501 | 0.008 |
| VWC | $CO_2$ | 0.488 | 0.010 |
| $N_{org}$ | $\delta^{13}C$ | 0.460 | 0.016 |
| $CO_2$ | $C_{org}$ | 0.449 | 0.019 |
| $CO_2$ | C/N | 0.446 | 0.020 |
| $CO_2$ | $N_{org}$ | 0.394 | 0.042 |
| $N_{org}$ | C/N | 0.388 | 0.046 |
| $CO_2$ | $\delta^{13}C$ | 0.386 | 0.047 |
| VWC | $N_{org}$ | 0.382 | 0.049 |
| GWC | $\delta^{13}C$ | 0.366 | 0.061 |
| GWC | C/N | 0.363 | 0.063 |
| VWC | $\delta^{13}C$ | 0.353 | 0.071 |
| VWC | $CH_4$ | -0.239 | 0.229 |
| $CH_4$ | $\delta^{15}N$ | -0.235 | 0.239 |
| $CO_2$ | $\delta^{15}N$ | -0.228 | 0.252 |
| $CH_4$ | GWC | 0.198 | 0.321 |
| $CH_4$ | $C_{org}$ | -0.194 | 0.333 |

| | | | |
|---|---|---|---|
| $\delta^{15}N$ | $\delta^{13}C$ | 0.153 | 0.446 |
| $CO_2$ | GWC | 0.108 | 0.590 |
| $CH_4$ | $N_{org}$ | -0.104 | 0.605 |
| $N_{org}$ | $\delta^{15}N$ | -0.096 | 0.633 |
| C/N | $\delta^{15}N$ | 0.084 | 0.679 |
| VWC | $\delta^{15}N$ | -0.067 | 0.739 |
| GWC | $\delta^{15}N$ | -0.064 | 0.752 |
| $C_{org}$ | $\delta^{15}N$ | -0.023 | 0.911 |

**Table S3:** Loading scores of environmental variables included in the initial PCA analysis. Included variables were: volumetric water content (VWC), gravimetric water content (GWC), $CH_4$, $CO_2$, $C_{org}$, $N_{org}$, $C_{org}/N_{org}$ ration, $\delta^{15}N$, $\delta^{13}C$.

| Parameter | PC1 | PC2 |
|---|---|---|
| VWC | -0.391 | -0.141 |
| $CH_4$ | 0.121 | 0.636 |
| $CO_2$ | -0.316 | -0.198 |
| GWC | -0.382 | 0.300 |
| $N_{org}$ | -0.426 | 0.265 |
| $C_{org}$ | -0.477 | 0.087 |
| C/N | -0.332 | -0.431 |
| $\delta^{15}N$ | -0.004 | -0.328 |
| $\delta^{13}C$ | 0.258 | -0.273 |

**Table S4:** Loading scores of environmental variables included in the final PCA analysis. Included variables were: volumetric gravimetric water content (GWC), $CH_4$, $CO_2$, $C_{org}$.

| Parameter | PC1 | PC2 |
|---|---|---|
| $CH_4$ | 0.157 | 0.787 |
| $CO_2$ | -0.503 | -0.383 |
| GWC | -0.547 | 0.476 |
| $C_{org}$ | -0.651 | 0.087 |

**Table S5:** peaks of qPCR values related to the $CH_4$ and nitrogen cycles, in the four sampled environments. For each environment, the peak expression is presented along with the corresponding depth. Samples were characterized based on sampling environments: SHE = Summer Shallow High Elevation (n=6), SLE = Summer Shallow Low Elevation (n=10), WLE = Winter Low Elevation (n=11). The WLE samples were comprised of shallow (<3 m, n=5) and deep (<3 m, n=6) samples. For elaboration, see materials and methods. p-value considered significant if < 0.05, and marked bold with underline.

| **Gene** | **Process** | **SHE peak** | **SLE peak** | **WLE peak Shallow samples (<3 m)** | **WLE peak Deep samples (>3 m)** |
|---|---|---|---|---|---|
| | | **Expression (depth) (copies/1gr soil)** | **Expression (depth) (copies/1gr soil)** | **Expression (depth) (copies/1gr soil)** | **Expression (depth) (copies/1gr soil)** |
| **mcrA** | **Methanogenesis** | $1.9x10^6 \pm 4.7x10^5$ (at 225 cm) | $5.3x10^6 \pm 3.5x10^5$ (at 166 cm) | $2.8x10^6 \pm 3.6x10^5$ (at 285 cm) | $2x10^6 \pm 4.9x10^5$ (at 711 cm) |
| **pmoA** | **Aerobic $CH_4$ oxidation** | $15.6x10^7 \pm 10.6x10^6$ (at 8 cm) | $4.05x10^7 \pm 1.02x10^6$ (at 68 cm) | $1.4\ x10^6 \pm 3.3x10^5$ (at 100 cm) | $1.4x10^5 \pm 1.8x10^4$ (at 711 cm) |
| **narG** | **Denitrification** | $3.9x10^7 \pm 7.5x10^5$ (at 8 cm) | $3.9x10^7 \pm 2x10^6$ (at 86 cm) | $3.4x10^6 \pm 1.6x10^5$ (at 100 cm) | $2.5x10^6 \pm 6.6x10^4$ (at 345 cm) |
| **nirK** | **Denitrification** | $3.3x10^6 \pm 4.5x10^4$ (at 8 cm) | $1.7x10^6 \pm 8.6x10^4$ (at 86 cm) | $1.1x10^6 \pm 5.2x10^4$ (at 162 cm) | $1.4x10^5 \pm 2x10^4$ (at 711 cm) |
| **norB** | **Denitrification** | 0 | $2.3\ x10^5 \pm 1.5x10^4$ (at 86 cm depth) | 0 | 0 |
| **hzsB** | **Anammox** | $2.1x10^6 \pm 6.8x10^4$ (at 8 cm) | $6.8x10^6 \pm 1.5x10^5$ (at 86 cm) | $2x10^6 \pm 1.8x10^3$ (at 162 cm) | $3.2x10^5 \pm 3x10^4$ (at 711 cm) |
| **A6-acm (16S)** | **Feammox** | $4.3x10^7 \pm 1.5x10^6$ (at 8 cm) | $2.9x10^7 \pm 6.8x10^5$ (at 86 cm) | $2.3x10^6 \pm 1.4x10^5$ (at 162 cm) | $3.2x10^5 \pm 2x10^4$ (at 535 cm) |
| **NC10 (16S)** | **NC10 phylum** | $6.5x10^4 \pm 4.2x10^3$ (at 8 cm) | $8.1x10^4 \pm 6.4x10^3$ (at 86 cm) | $1.4x10^5 \pm 1.4x10^4$ (at 100 cm) | 0 |
| **amoA archaea** | **Aerobic ammonium oxidation** | $3.6x10^5 \pm 9.4x10^3$ (at 8 cm) | $3.2x10^5 \pm 1.7x10^4$ (at 23 cm) | 0 | $7.3x10^4 \pm 9.2x10^3$ (at 711 cm) |
| **amoA bacteria** | | $1.7x10^6 \pm 3.8x10^4$ (at 8 cm) | $2.4x10^6 \pm 1x10^5$ (at 56 cm) | $3.6x10^5 \pm 1.9x10^4$ (at 162 cm) | $1.4x10^4 \pm 1.7x10^3$ (at 345 cm) |

**Table S6:** Kruskal-Wallis rank sum tests for 16S NGS alpha diversity between sampled environments: SHE = Summer Shallow High Elevation (n=6), SLE = Summer Shallow Low Elevation (n=10), WLE = Winter Low Elevation (n=11). The WLE samples were comprised of shallow (<3 m, n=5) and deep (<3 m, n=6) samples. For elaboration, see materials and methods. p-value considered significant if < 0.05, and marked bold with underline.

| Comparison | Alpha index | df | W statistic | p-value |
|---|---|---|---|---|
| **Sampling environment** | **Shannon’s entropy** | 3 | 5.41 | 0.144 |
| | **Faith's PD** | 3 | 19.18 | **0.0002** |
| | **Pielou's evenness** | 3 | 6.9 | **0.074** |

**Table S7:** Post hoc via Dunn's tests of 16S NGS alpha diversity between sampled environments: SHE = Summer Shallow High Elevation (n=6), SLE = Summer Shallow Low Elevation (n=10), WLE = Winter Low Elevation (n=11). The WLE samples were comprised of shallow (WSLE, <3 m, n=5) and deep (WDLE, <3 m, n=6) samples. p-value considered significant if < 0.05, and marked bold with underline.

| Alpha index | group1 | group2 | n1 | n2 | statistic | p-value | p.adj |
|---|---|---|---|---|---|---|---|
| Faith pd | SHE | SLE | 6 | 10 | -0.195 | 0.845 | 0.845 |
| | SHE | WDLE | 6 | 5 | -2.800 | 0.005 | **0.008** |
| | SHE | WSLE | 6 | 6 | -3.204 | 0.001 | **0.004** |
| | SLE | WDLE | 10 | 5 | -2.936 | 0.003 | **0.007** |
| | SLE | WSLE | 10 | 6 | -3.358 | 0.001 | **0.004** |
| | WDLE | WSLE | 5 | 6 | -0.534 | 0.593 | 0.712 |

**Table S8:** PERMANOVA and PERMDISP results of 16S NGS beta diversity. Samples were grouped based on sampling environments: SHE = Summer Shallow High Elevation (n=6), SLE = Summer Shallow Low Elevation (n=10), WLE = Winter Low Elevation (n=11). The WLE samples were comprised of shallow (WSLE, <3 m, n=5) and deep (WDLE, <3 m, n=6) samples. For elaboration, see materials and methods. Number of permutations = 999. p-value considered significant if < 0.05, and marked bold with underline.

| Beta index | Test | Comparison | n | N (groups) | test statistic | p-value |
|---|---|---|---|---|---|---|
| Jaccard | PERMDISP | Sampling environment | 27 | 4 | 1.73 | 0.264 |
| Bray Curtis | PERMDISP | Sampling environment | 27 | 4 | 1.188 | 0.428 |
| Jaccard | PERMANOVA | Sampling environment | 27 | 4 | 2.402 | **0.001** |
| Bray Curtis | PERMANOVA | Sampling environment | 27 | 4 | 3.798 | **0.001** |

**Table S9:** PERMANOVA and PERMDISP post hoc pairwise comparison results of 16S NGS beta diversity. Samples were grouped based on sampling environments: SHE = Summer Shallow High Elevation (n=6), SLE = Summer Shallow Low Elevation (n=10), WLE = Winter Low Elevation (n=11). The WLE samples were comprised of shallow (WSLE, <3 m, n=5) and deep (WDLE, <3 m, n=6) samples. For elaboration, see materials and methods. Number of permutations = 999. p-value considered significant if < 0.05, and marked bold with underline.

| Bata index | Test | Comparison | Group 1 | Group 2 | n | pseudo-F | p-value | q-value |
|---|---|---|---|---|---|---|---|---|
| **Jaccard** | **PERMANOVA** | **Sampling environment** | SHE | SLE | 16 | 999 | 1.787432178 | **0.018** |
| | | | SHE | WDLE | 11 | 999 | 3.311730722 | **0.003** |
| | | | SHE | WSLE | 12 | 999 | 1.98687807 | **0.003** |
| | | | SLE | WDLE | 15 | 999 | 3.271207712 | **0.001** |
| | | | SLE | WSLE | 16 | 999 | 1.741631202 | **0.008** |
| | | | WDLE | WSLE | 11 | 999 | 2.568510894 | **0.001** |
| **Bray Curtis** | **PERMANOVA** | **Sampling environment** | SHE | SLE | 16 | 999 | 2.658819688 | **0.002** |
| | | | SHE | WDLE | 11 | 999 | 4.501196181 | **0.002** |
| | | | SHE | WSLE | 12 | 999 | 2.539094434 | **0.007** |
| | | | SLE | WDLE | 15 | 999 | 6.366077494 | **0.001** |
| | | | SLE | WSLE | 16 | 999 | 3.422828237 | **0.001** |
| | | | WDLE | WSLE | 11 | 999 | 3.221380856 | **0.002** |

**Table S10:** ADONIS test results of 16S NGS beta diversity with selected environmental variables. Number of permutations = 999. MGC = Gravimetric Water Content. Variables were chosen based on the principal component analysis (Fig. 3A-B and Table S4). p-value considered significant if < 0.05, and marked bold with underline.

| Beta index | Variable | df | SumsOfSqs | MeanSqs | F.Model | R2 | Pr(>F) |
|---|---|---|---|---|---|---|---|
| **Jaccard** | **GWC** | 1 | 0.652 | 0.652 | 1.951 | 0.060 | **<u>0.003</u>** |
| | **ch4** | 1 | 0.910 | 0.910 | 2.724 | 0.084 | **<u>0.001</u>** |
| | **co2** | 1 | 0.445 | 0.445 | 1.331 | 0.041 | 0.060 |
| | **Corg** | 1 | 0.364 | 0.364 | 1.089 | 0.034 | 0.273 |
| | **env** | 3 | 2.104 | 0.701 | 2.099 | 0.194 | **<u>0.001</u>** |
| | **Residuals** | 19 | 6.349 | 0.334 | NA | 0.587 | NA |
| | **Total** | 26 | 10.823 | NA | NA | 1.000 | NA |
| **Bray Curtis** | GWC | 1 | 0.839 | 0.839 | 3.398 | 0.088 | **<u>0.001</u>** |
| | ch4 | 1 | 0.979 | 0.979 | 3.964 | 0.102 | **<u>0.001</u>** |
| | co2 | 1 | 0.364 | 0.364 | 1.476 | 0.038 | 0.091 |
| | Corg | 1 | 0.281 | 0.281 | 1.138 | 0.029 | 0.291 |
| | env | 3 | 2.422 | 0.807 | 3.271 | 0.253 | **<u>0.001</u>** |
| | Residuals | 19 | 4.690 | 0.247 | NA | 0.490 | NA |
| | Total | 26 | 9.575 | NA | NA | 1.000 | NA |

**Table S11**: Primers and gBlocks used in this study. gBlocks were constructed based on the sequences of relevant gene, denoted by their NCBI accession number. The full references are presented at the main manuscript.

| **Gene** | **Biogeochemical Function** | **Primer name** | **Sequence (5'-3')** | **Length (bp)** | **Primer concentration (mM)** | *** gBlocks NCBI accession number** | **References** |
|---|---|---|---|---|---|---|---|
| mcrA | methanogenesis | ME1F<br>ME3R | GCMATGCARATHGGWATGTC<br>TGTGTGAASCCKACDCCACC | 350 | 0.5 | NC_014507.1 | [107, 108] |
| pmoA | $CH_4$ oxidation | A189gc<br>mb661 | GGNGACTGGGACTTCTGG<br>CCGGMGCAACGTCYTTACC | 472 | 0.5 | L40804.2 | [109–111] |
| nirK | denitrification | F1aCu<br>R3Cu | ATCATGGTSCTGCCGCG<br>GCCTCGATCAGRTTGTGGTT | 473 | 0.15 | EF623493.1 | [112] |
| narG | denitrification | narG571F<br>narG773R | CCGATYCCGGCVATGTCSAT<br>GGNACGTTNGADCCCCA | 203 | 0.5 | Based on plasmid | [113] |
| norB | denitrification | qnorB2<br>qnorB5R | GGNCAYCARGGNTAYGA<br>ACCCANAGRTGNACNACCCACCA | 263 | 0.5 | Based on plasmid | [114] |
| amoA bacteria | Aerobic ammonium oxidation | amoA-1F<br>amoA-2R | GGGGTTTCTACTGGTGGT<br>CCCCTCKGSAAAGCCTTCTTC | 429 | 0.5 | U76552.1 | [115, 116] |
| amoA archaea | Aerobic ammonium oxidation | Arch-amoA-F<br>Arch-amoA-R | STAATGGTCTGGCTTAGACG<br>GCGGCCATCCATCTGTATGT | 635 | 0.25 | MF176967.1 | [117, 118] |
| hzsB | ANAMMOX | HSBeta396F<br>HSBeta742R | ARGGHTGGGGHAGYTGGAAG<br>GTYCCHACRTCATGVGTCTG | 385 | 0.5 | KP002830.1 | [119, 120] |
| 16S | Feammox (Acidimicrobiaceae sp. strain A6) | acm342f<br>acm439r | GCAATGGGGGAAACCCTGAC<br>ACCGTCAATTTCGTCCCTGC | 200 | 0.15 | MG589453.1 | [121] |
| 16S | NC10 phylum | qp1F<br>qp1R | GGGCTTGACATCCCACGAACCTG<br>CGCCTTCCTCCAGCTTGACGC | 203 | 0.15 | MZ778768.1 | [122] |
