## Supplementary material for "Complex nitrogen redox couplings control methane emissions from Arctic upland yedoma taliks": Description of Additional Supplementary Files

**Title: Supplementary Data 1**

**Description: 16S ASV table.** For each ASV, the full MD5 code (Hash_ID) is presented. Additional information provided is occurrence per sample (counts), QIIME2 classification, representative sequence and confidence. For each sample we included the identity of sampling environment, core and depth (in cm).

Sampling environment: **SSHE** = Summer Shallow High Elevation (n=6), **SSLE** = Summer Shallow Low Elevation (n=10), **WSLE** = Winter Shallow Low Elevation (n=6), **WDLE** = Winter Deep Low Elevation (n=5).

**Title: Supplementary Data 2**

**Description: 16S genus level abundance.** ASVs were collapsed at the genus level, utilizing the QIIME2 qiime taxa collapse function. Additional information provided is occurrence per sample (counts), QIIME2 classification, representative sequence and confidence. For each sample we included the identity of sampling environment, core and depth (in cm). Bacteria and Archaea total counts and relative abundances are also presented.

Sampling environment: **SSHE** = Summer Shallow High Elevation (n=6), **SSLE** = Summer Shallow Low Elevation (n=10), **WSLE** = Winter Shallow Low Elevation (n=6), **WDLE** = Winter Deep Low Elevation (n=5).

**Title: Supplementary Data 3**

**Description: 16S archaea phyla relative abundance.** ASVs for archaea only were collapsed at the phylum level, utilizing the QIIME2 qiime taxa collapse function. Additional information provided is occurrence per sample and per sampling environment (relative abundance), and QIIME2 phyla classification. For each sample we included identity of sampling environment, core and depth (in cm).

Sampling environment: **SSHE** = Summer Shallow High Elevation (n=6), **SSLE** = Summer Shallow Low Elevation (n=10), **WSLE** = Winter Shallow Low Elevation (n=6), **WDLE** = Winter Deep Low Elevation (n=5).

**Title: Supplementary Data 4**

**Description: 16S bacterial phyla relative abundance.** ASVs for bacteria only were collapsed at the phylum level, utilizing the QIIME2 qiime taxa collapse function. Additional information provided is occurrence per sample and per sampling environment (relative abundance), and QIIME2 phyla classification. For each sample we included identity of sampling environment, core and depth (in cm).

Sampling environment: **SSHE** = Summer Shallow High Elevation (n=6), **SSLE** = Summer Shallow Low Elevation (n=10), **WSLE** = Winter Shallow Low Elevation (n=6), **WDLE** = Winter Deep Low Elevation (n=5).

**Title: Supplementary Data 5**

**Description: 16S rRNA gene table of methanogens and methanotrophs obtained by next generation sequencing (NGS).** Full taxonomy is presented, occurrence per sample and per sampling environment (relative abundance). For each sample we included identity of sampling environment, core and depth (in cm).

Sampling environment: **SSHE** = Summer Shallow High Elevation (n=6), **SSLE** = Summer Shallow Low Elevation (n=10), **WSLE** = Winter Shallow Low Elevation (n=6), **WDLE** = Winter Deep Low Elevation (n=5).

**Title: Supplementary Data 6**

**Description: PICRUST2 KEGG ORTHOLOGY, based on 16S rRNA gene NGS data.** Data is presented as predicted functional abundances for all identified pathways.

**Title: Supplementary Data 7**

**Description: PICRUST2 KEGG modules, based on 16S rRNA gene NGS data.** Data is presented as relative abundance for all identified modules.
